# Chronic IL-1-induced DNA double-strand break response in hippocampal neurons drives cognitive deficits

**DOI:** 10.1101/2024.02.03.578747

**Authors:** Belloy Marcy, Benjamin A. M. Schmitt, Florent H. Marty, Paut Charlotte, Bassot-Parra Emilie, Aïda Amel, Alis Marine, Ecalard Romain, Boursereau Raphaël, Gonzalez-Dunia Daniel, Blanchard Nicolas, Suberbielle Elsa

**Affiliations:** Infinity, Université Toulouse, CNRS, Inserm, UPS, Toulouse, France; Inserm US006, ANEXPLO/CREFRE, Purpan Hospital, Toulouse, France

## Abstract

Chronic inflammation characterized by increased cytokine levels, such as interleukin-1 (IL-1), accompanies many neurological diseases but little is known about IL-1 contribution to cognitive impairment and its interplay with epigenetic processes, including the DNA double-strand break (DSB) response.

Here, we demonstrate that H2A.X-dependent DSB signaling in hippocampal neurons drives cognitive deficits upon chronically elevated IL-1. Mice persistently and latently infected with *Toxoplasma gondii* display impaired spatial memory consolidation along with elevated IL-1β in the hippocampus. We find that neuronal IL-1 signaling in excitatory neurons is required for the spatial memory deficits caused by *T. gondii* infection and by chronic systemic infusion of IL-1β. In both cases, the deficit in spatial memory was prevented by the abrogation of neuronal H2A.X-dependent signaling.

Our results highlight the instrumental role of cytokine-induced DSB-dependent signaling in spatial memory defects. This novel pathological mechanism in inflammation control of neuronal function may extend to several neurological diseases.

## INTRODUCTION

Chronic neurological diseases are devastating pathologies with a major medical and economic burden. Although their underlying etiologies and molecular mechanisms are not fully understood, the neuronal inflammatory response has been identified as a key player in neurological dysfunction.

Chronic neuroinflammation may be sterile or caused by the invasion and persistence of microorganisms in the central nervous system (CNS), such as viruses or parasites ^1^. Although such infections may seem mostly asymptomatic, the persistence of a pathogen in the CNS substantially shapes the local innate and adaptive immune responses, with potential long-term consequences on brain function such as inflammaging and even neurodegeneration ^2–4^.

For pathogens that establish persistent infections in the CNS, including Herpesviruses, Bornavirus, and *Toxoplasma gondii (T. gondii* or *Tg)*, intricate relationships between host neural cells, immune responses and the pathogens are determinant for disease outcome. *T. gondii* is an intracellular apicomplexan parasite that can establish long-lasting CNS infection in all warm-blooded animals, including humans. Thirty (30) to 50% of the worldwide population is estimated to display a positive serology for (i.e. has been exposed to) *T. gondii* ^5^. Although this infection can lead to a deadly encephalitis in immunocompromised patients due to parasite reactivation in the brain, most cases of infection in immunocompetent subjects are seemingly clinically silent thanks to effective immune control of the parasite during the acute and chronic stages, and hence are called “latent”. Nonetheless, a substantial body of evidence suggests that latent *T. gondii* infection contributes to several human neuropsychiatric disorders such as schizophrenia, bipolar disorder, obsessive compulsive disorder and epilepsy ^6–10^. Studies of *T. gondii* experimental infection in the mouse, one of the parasite’s natural hosts, have indicated that the parasite burden in the brain, the level of neuroinflammation, and the extent of neurological symptoms vary with the combination of parasite strain and mouse genetic background, most likely since these processes are influenced by the efficiency of parasite immune control in the brain. Several (neuro)immune cells contribute to *T. gondii* immune surveillance in the CNS, including microglia ^11^, astrocytes ^12,13^, monocytes ^14^, and CD8+ T lymphocytes, which play a central role ^15,16^. C57BL/6 mice are known to be naturally susceptible to develop encephalitis upon infection by type II *T. gondii.* However, infection of C57BL/6 mice with type II parasites which are genetically engineered to express a model antigen that is efficiently processed and presented by major histocompatibility class I (MHCI) molecules, elicits protective CD8+ T cell responses which provide effective brain parasite control, and allow the establishment of latency ^17^. So far, most behavioral and cognitive impairments have been studied in the context of *T. gondii*-induced chronic encephalitis. Exploration of the contribution of latent infection to behavioral alterations remains mostly limited to innate and anxiety-related behavior, whereas the impact on more complex cognitive functions (e.g. memory) is ill-defined ^18,19^. Therefore, studies are needed to understand the neural processes that may be impacted by latent *T. gondii* infection, and that may cause cognitive dysfunction. Interestingly, recent evidence have suggested a detrimental role for neuroinflammation in *T. gondii*-induced behavioral alterations ^19–21^. However, a full understanding of the causes of neuronal dysfunction upon infection is challenging, due to the intricate intervention of inflammatory and neuronal components in the pathology.

The ability of the immune response to control infection relies in great part on the production of cytokines. Cytokines are known to play both beneficial and detrimental roles in the CNS ^22–26^. While basal signaling by Interferon-γ ^27^, Interleukin (IL)-17 ^28^, IL-4 ^29^, IL-1α ^30^, IL-33 ^31^ play important neuromodulatory roles underlying social behavior, short-term or fear memory encoding, respectively, IL-1β, IL6, IL-17, IL-33 ^32,33^, Tumor Necrosis Factor-α, and Interferon α and γ also mediate chronic pathological signaling, which in turn leads to neuronal dysfunction ^22,34^. Chronically elevated levels of circulating IL-1, IL-6, TNF-α, or IFN-α lead to persistent alterations in neurotransmitters like glutamate, and impair neuronal growth factor function ^25,35^. Thus, although the molecular mechanisms underlying the conversion of cytokine signals into neuro-pathological factors have been partly described ^22^, the mechanisms by which they exert a direct and long-lasting impact on neuronal function require further investigation.

One hypothesis to explain the long-lasting nature of behavioral alterations, is that inflammation could perturb epigenetic mechanisms. These mechanisms are known to be key regulators of neuronal function. Thanks to tightly controlled sequences of molecular events, they allow at the same time a highly dynamic response adjusting to the environment and a durable impact on gene expression by maintenance of epigenetic marks ^36^. Indeed, indirect evidence suggests that pro-inflammatory cytokines modulate the epigenome of neural cells ^37–39^. DNA double-strand breaks (DSB), and their associated response, drive complex epigenetic processes^40–43^ and are now considered as a central regulator of cognition ^40–43^. In neurodegenerative conditions such as Alzheimer’s disease (AD), it was shown that DSB and one of their markers, the phosphorylated histone variant H2A.X, accumulate in neurons, because of impaired DSB repair mechanisms due to disturbed electrical activity of neuronal networks in the brain, leading to cognitive decline ^41,42^. Interestingly, similar alterations in electrical activity are also detected during exposure to pro-inflammatory cytokines ^22,44,45^. In this study, we hypothesized that, due to its inflammatory component, latent *T. gondii* infection may cause cognitive deficits and that their underlying mechanisms may converge onto alterations in the neuronal response to DSB. Using mouse models, we showed that chronic infection by *T. gondii* impairs specific components of hippocampus-dependent memory. These impairments could be mimicked by chronic exposure to low blood concentrations of IL-1β. They required activation of the IL-1-dependent signaling pathway in glutamatergic neurons, and could be prevented by invalidating the H2A.X-dependent DNA DSB response in neurons. Together, our data show that a persistent, low-grade increase in IL-1β level, whether it is local (CNS) or systemic, is sufficient to cause precision memory deficits by inducing an excessive DNA DSB response.

## RESULTS

### Chronic *Toxoplasma gondii* infection impairs spatial consolidation and precision memory

First, to fully characterize the impact of *T. gondii* infection on cognitive behaviors, we performed a battery of behavioral tests on mice infected with either of two genetically-engineered *T. gondii* lines derived from the same type II Prugniaud strain: the *T. gondii*.SAG1-OVA line, which causes toxoplasmic encephalitis in mice of C57BL/6J background, and the *T. gondii*.GRA6-OVA line, which leads to latent infection characterized by lower brain parasite load (Extended Data Fig.1A, B) due to effective CD8+ T cell-mediated parasite immune surveillance in the CNS ^17^. Behavioral testing was performed once all infected mice had recovered from the acute phase of the infection and reached back 90% of their initial weight (Extended Data Fig.1C). To test the learning and memory capacities of the mice upon *T. gondii* infection, mice were challenged in the Barnes maze task. In the training phase of the Barnes maze, all groups were able to learn the task and improved over time, since they all ran shorter distances and committed less and less errors over time prior to finding the target hole (Fig. 1A,B). Yet, mice with encephalitis were slower to improve their performances as they visited more error holes (Fig. 1B) and adopted less efficient, non-spatial, search strategies of the target hole (Fig. 1C,D). Nonetheless, after 5 days of training, all groups had reached similar performances, allowing us to test the recall of spatial memory in a probe trial set 24h after the last training (Fig. 1E,F). Mice from all groups displayed preserved approximate memory of the target hole, since they spent significantly more time in the target quadrant where the exit hole used to be, compared to the chance threshold (Fig. 1E). However, during the probe test, mice with encephalitis displayed less precision in visiting the area as indicated by fewer visits of the exit area compared to uninfected mice (Fig. 1F). Interestingly, latently infected mice also visited significantly less the exit area than uninfected mice (Fig. 1F), suggesting discrete impairment in memory retrieval upon both encephalitis and latent *T. gondii* infection. In order to better dissect the regional network mechanisms involved in these impairments that are common to the latency and encephalitis contexts, we performed two object-based tests: the novel object recognition test (NOR, Fig. 1G,H) and the object location test (OL, Fig. 1I,J). In the experimental conditions used for these tests, the NOR assesses memory retrieval and involves cortical areas of the brain, whereas the OL task evaluates the consolidation of spatial memory and critically relies upon the function of the hippocampus. Mice from all groups spent the same amount of time exploring objects (Fig. 1G & 1I), and properly discriminated a new object compared to an already explored one, suggesting intact cortical circuits despite *T. gondii* infection (Fig. 1H). Yet, *T. gondii*-infected mice, regardless of disease severity, showed impaired consolidation of spatial memory since they were unable to discriminate the moved object (Fig. 1J). Together, these data indicate that *T. gondii*-infected mice with either encephalitis or latency are impaired in spatial memory retrieval and consolidation, which both depend on the function of the hippocampus.

**Figure 1:**
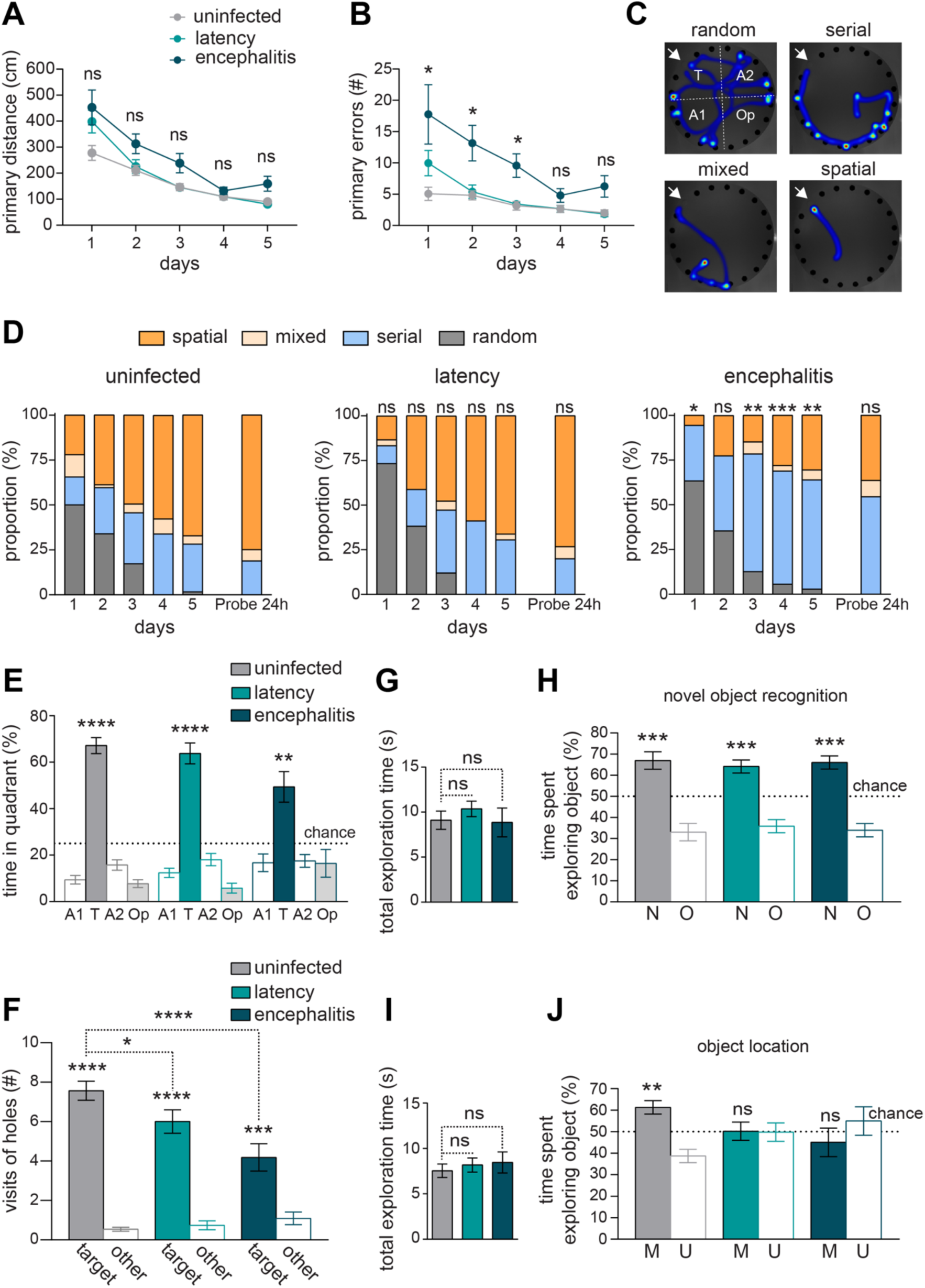
*Toxoplasma gondii* chronic infection impairs the precision and consolidation of spatial memory in a severity-dependent manner. Starting 12 weeks post-infection by *Tg*.GRA6-OVA (latency) *vs*. *Tg*.SAG1-OVA (encephalitis) parasites, or PBS (uninfected), C57BL6/J mice were tested in a series of behavioral tests: the Barnes maze (**A-F**), the novel object recognition test (**G-H**) and the object location task (**I-J**). **A-B**) In the Barnes maze, learning curves show mean daily distance run (**A**), or mean numbers of errors of holes visits (**B**) prior to find the target hole connected to a hidden exit box. Effect of the time (p<0.0001) and *Tg* strain (p<0.001) with no interaction as analyzed by repeated measures mixed-effects models, and Dunnett pos-hoc tests compared to the non-infected group. (**C-D**) The strategy used by mice to find the target hole was determined. **C**) Representative trajectory heatmaps illustrating the four types of strategies used to find the target hole indicated by the white arrow. White dashed lines delineate quadrants. A1: adjacent quadrant 1, T: target quadrant, A2: adjacent quadrant 2, Op: opposite quadrant **D**) Mean distribution of daily strategies in the course of the 6 training days and the probe trial, as shown as percentage per group. Freeman-Halton extension of the Fisher ’s exact test using 3 groups (mixed and spatial groups were merged) was used to compare distributions with uninfected mice. **E**) Percentage of time that mice spent in the target quadrant *vs*. the three other quadrants during a 90-s probe trial, run 24 h after the last training trial in the Barnes. Results were tested by the one-sample t-test compared to chance (25%). **F**) Number of visits of the original exit hole (target) location *vs*. each other hole location in other quadrants (other) during the probe trial. Results were analyzed by paired *t*-test or by Dunnett post-hoc tests compared to the non-infected group as indicated by brackets. *n* = 11–16 mice per group of treatment pooled from 2 independent cohorts. (**G-H**) General long-term memory assessed in a novel object recognition task, 24 h after the first exploration. Bars in **H**) represent the time spent by mice exploring the object that is novel (New, N) or old (Old, O), expressed as percentage out of the total time spent exploring either object (shown in **G**). (**I-J**) Consolidation of spatial memory was assessed in the object location task, 3 h after initial exploration. Bars in **J**) represent the time spent by mice exploring the object that has been moved (M) or not (unmoved, U), expressed as percentage out of the total time spent exploring either object (shown in I). (**G-J**) *n* = 10–20 mice per group of treatment pooled from 2 independent cohorts. Results were tested in **G-J** by one-sample t-test compared to chance (50%). ns, non-significant, *p<0.05, **p<0.01, ***p<0.001, ****p<0.0001. Values are indicated as means ± s.e.m.

### Chronic *Toxoplasma gondii* infection upregulates IL-1 dependent pathways in the hippocampus, which are responsible for the cognitive impairment

To start investigating the cellular mechanisms which may lead to impaired consolidation of spatial memory, we assessed the changes in numbers and activation state of non-neuronal cell populations in the hippocampus (e.g. astrocytes, microglia, monocytes, granulocytes and T cells) by immunohistofluorescence (Extended Data Fig. 1 D,E) and flow cytometry (Extended Data Fig. 1 F-O). The number of astrocytes, microglia, monocytes, neutrophils/granulocytes, Ly6C^lo^ monocyte-derived cells and T cells, as well as the activation of microglia (measured by MHC class II and CD86 surface expression levels), were all increased in the hippocampus of infected mice compared to uninfected controls (Extended Data Fig. 1 D-O). Despite a clear trend for a larger increase in the encephalitic mice, the differences between the encephalitis and latency conditions did not reach statistical significance. Thus, both models of chronic *T. gondii* persistence in the brain lead to peripheral immune cell infiltration, astrogliosis, microgliosis, and microglia activation.

To gain insights into the cytokine-related pathways that are dysregulated during chronic *T. gondii* infection and may cause behavioral abnormalities, we analyzed gene expression changes in the hippocampus of uninfected *vs*. infected mice by bulk RNA-sequencing, and we used these data to infer pathways significantly enriched in both latency and encephalitic mice *vs*. uninfected mice. When filtering for significantly enriched pathways related to all interleukins or interferons, we found pathways linked to production and response to IFN-γ, type I IFN, IL-1, IL-10 and IL-27 (Fig. 2A,B). Given i) that IL-1 ^30,46^ and IL-1-related cytokines (e.g. IL-33 ^33,47^) have been suggested to modulate hippocampal function, ii) that the IL-1 receptor is constitutively expressed in hippocampal neurons of the dentate gyrus ^48^ and iii) that IL-1 signaling does not seem to be required for parasite control during chronic stage ^49^, we decided to further explore the role of brain IL-1-signaling in latent *T. gondii*-induced behavioral abnormalities.

**Figure 2:**
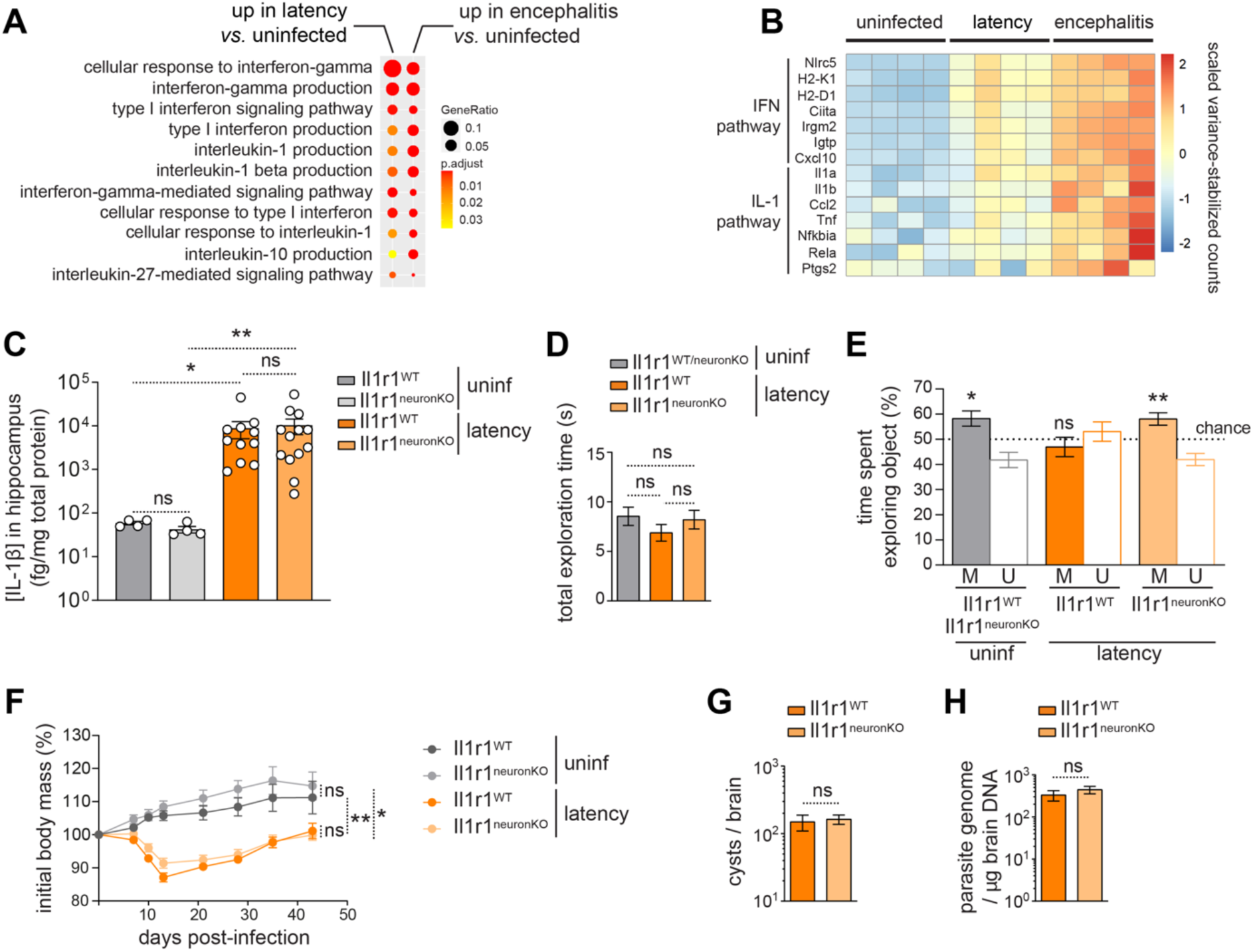
Neuronal IL-1-dependent signaling pathways drive cognitive impairment caused by *Tg* chronic infection. (**A-B**) Bulk RNA-sequencing analysis was performed on RNA prepared from hippocampi of mice chronically infected by *Tg*, either in the latency or encephalitis conditions versus non infected mice. **A**) Dot plot representation of pathway enrichment analyses focused on interferon and interleukin pathways that are significantly enriched in the infected groups compared to non-infected controls. Each dot indicates the magnitude of the changes (size of dot), and the statistical significance of the changes (color of dot). **B**) Heatmap showing the scaled variance-stabilized counts of 7 emblematic genes among the IFN and IL-1 pathways in uninfected, latency and encephalitis groups. Each square represents the data from one mouse. n=4 mice per group from 1 single experiment. (**C-H**) Consolidation of spatial memory was assessed in the object location task, 3 h after the initial exploration of *Tg*-infected *vs*. non infected transgenic mice, in which the IL-1R1 receptor was knocked out in glutamatergic excitatory neurons of the forebrain (*Il1r1^neuronKO^*) upon tamoxifen treatment 1 month prior to infection. (**C**) IL-1β concentration in mouse hippocampus at 7 wpi (Kruskal-Wallis, Dunn’s multiple comparison test, n = 4-13 mice per group from one experiment) Each dot represents one individual value. (**D-E**) Object location task: Bars in **E**) represent the time spent by mice exploring the object that was moved (M) or not (unmoved, U), expressed as percentage out of the total time spent exploring either object (shown in **D**). n = 11-14 mice per group of treatment pooled from 2 independent cohorts (one-sample *t* test compared to chance at 50 %). (**F**) Percentage of initial body mass throughout infection, n = 4-13 mice per group from one experiment (Kruskal-Wallis and Dunn’s multiple comparison test performed on the area under the curve). (**G**) Cyst number and (**H**) total parasite burden in the brain, n= 11-13 infected mice per group from one experiment (unpaired Student’s *t* test). ns, non-significant, *p<0.05, **p<0.01. Graphs show means ± s.e.m.

To this aim, we implemented a transgenic mouse model, wherein IL-1-dependent signaling can be suppressed in adult excitatory neurons by conditional CaMKIIα-restricted ^50^ and tamoxifen-inducible knockout of the receptor for IL-1 (IL-1R1) ^51^.

We confirmed that the IL-1β cytokine is more abundant in hippocampal protein extracts from latently infected compared to uninfected mice, and we observed that the absence of IL-1R1 in neurons did not change this level (Fig. 2C).

Despite the loss of IL-1 signaling in excitatory neurons, uninfected *Il1r1*^neuronKO^ mice displayed normal weight evolution (Extended Data Fig. 2A) normoactive and normo-anxious behavior (Extended Data Fig. 2B,C), and similar capacities of learning, consolidating and retrieval of memory compared to tamoxifen-treated Cre-negative control littermates (Extended Data Fig. 2D-H)). Since latent *T. gondii* infection causes impaired consolidation of spatial memory in the OL test, we tested *Il1r1*^WT^ and *Il1r1*^neuronKO^ mice chronically infected by latent *T. gondii* in the same OL task (Fig. 2D,E). We confirmed that latent *T. gondii*-infected *Il1r1*^WT^ mice were impaired in the OL task compared to uninfected littermates. Importantly, *Il1r1*^neuronKO^ mice displayed intact spatial consolidation in the OL task (Fig. 2E). Also, weight loss and brain parasite burden were unaffected by this conditional knockout (Fig. 2F-H). Hence, suppressing IL-1R1 signaling in excitatory neurons is sufficient to prevent the deleterious impact of chronic *T. gondii* infection on consolidation of spatial memory. Together, our data suggest that IL-1 signaling in neurons critically contribute to the memory deficits caused by chronic *T. gondii* infection.

### Chronic systemic exposure to low levels of IL-1β impairs the precision and consolidation of spatial memory due to a direct effect on glutamatergic neurons

Because IL-1β can be detected in the blood and/or in the brain of many chronic neurological diseases involving low-grade neuroinflammation, such as depression, brain injury or bipolar disorders ^25,30,52–54^, or latent *T. gondii* infection (see Fig. 2C), we decided to explore whether the deleterious impact of latent *T. gondii* infection on cognitive processes could be mimicked using chronic systemic administration of low concentrations of IL-1β. In other words, we sought to test if in addition to being required, neuronal IL-1 signaling would be sufficient to impair spatial memory. To address this question, we chronically exposed wild-type mice to 5μg/kg/day of recombinant mouse IL-1β using subcutaneously implanted osmotic minipumps. These devices allow to raise the serum IL-1β levels for up to 5 weeks (Fig. 3A). A transient and modest weight loss was observed during the first 48 hours post-surgery, which was more pronounced in IL-1β-treated mice than saline controls (Extended Data Fig. 3A). IL-1β-treated mice were slightly slower to regain weight during the first 7 days post-surgery. However, by two weeks of infusion, mice were indistinguishable in terms of weight and general behavior. During the last two weeks of infusion, mice were subjected to the same battery of behavioral challenges as the *T. gondii*-infected mice (Fig. 3 and Extended Data Fig. 3). In the elevated plus maze test (EPM), IL-1β-treated mice appeared normoactive and normo-anxious (Extended Data Fig. 3B,C). In the training phase of the Barnes maze, mice from both groups displayed normal learning abilities and performed equally well in terms of distance and numbers of errors prior to finding the exit hole (Fig. 3B,C). The strategies deployed by all groups to find the exit hole also evolved from poorly efficient to spatial strategies over the 5 days of training (Fig. 3D). When challenged with a probe trial 24h after the last training, mice from both groups remembered the global area of the exit hole, since they spent significantly more time in the target quadrant during the trial (Fig. 3E). Interestingly, however, IL-1β-treated mice explored less precisely this area, as they performed statistically less visits of the target hole than saline-treated controls (Fig. 3F). As indicated by the strategies to find the target hole, although not statistically significant, IL-1β-treated mice tended to less strictly adopt the spatial strategy to identify the target hole during the probe trial (see Fig. 3D), suggesting that a component of spatial memory was impaired. In order to determine which component of the memory was impaired in IL-1β-treated mice, we performed, as for *T. gondii* infection, the NOR (Fig. 3G) and OL (Fig. 3H) tests. Mice from all groups were able to discriminate a new object compared to a previously explored object in the NOR (Fig. 3G). However, IL-1β-treated mice showed impaired consolidation of spatial memory since they were unable to discriminate the moved object in the OL test, and explored both objects equally (Fig. 3H). To examine the durability of these effects, mice were challenged in the Barnes maze paradigm one month after the end of IL-1β infusion (Extended Data Fig.3D-G). First, we confirmed that serum IL-1β levels had returned below the detection limit and were indistinguishable from the saline-treated levels (Extended Data Fig. 3D). Both groups were able to learn and progress in their ability to find the target hole over the 5 days of training (Extended Data Fig. 3E,F). Nonetheless, IL-1β-treated mice continued to perform less precisely, as they made significantly less visits of the target hole compared to saline-treated controls (Extended Data Fig. 3G). These results suggest that despite serum weaning from IL-1β, the effects of chronic IL-1β administration on memory are long-lasting.

**Figure 3:**
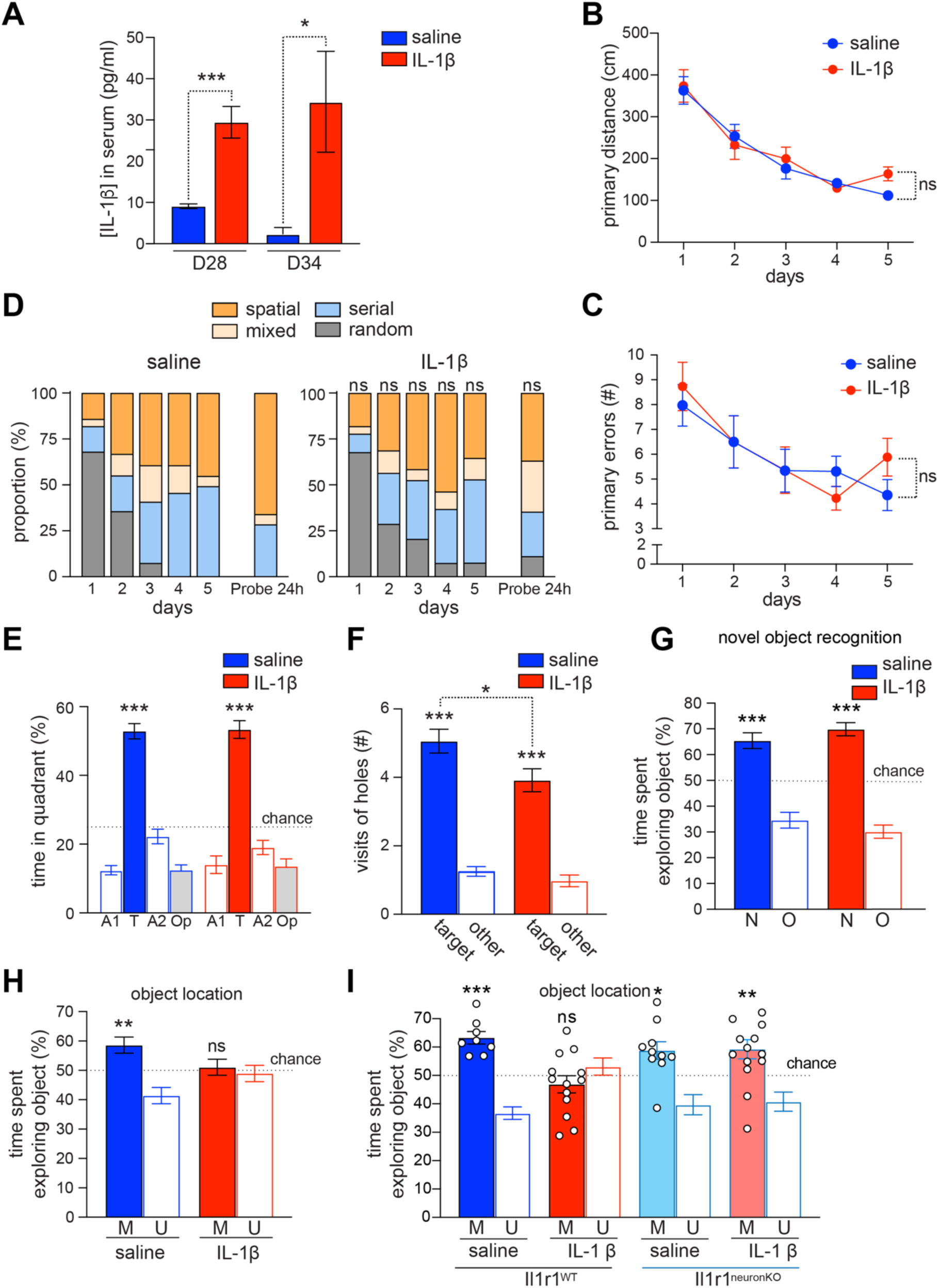
Chronic exposure to IL-1β induces deficits in the consolidation and the precision of spatial memory. C57BL/6J male mice were subcutaneously implanted with osmotic mini-pumps containing saline solution or IL-1β diluted in saline solution (5 µg/kg/day) for 28 days or 35 days. Starting 3 weeks post-implantation, mice were tested in a series of behavioral tests: the Barnes maze (B-F), the novel object recognition test (G) and the object location task (H-I). **A**) IL-1β concentrations in the serum were assessed by ELISA in saline controls and IL-1β-treated mice. n= 11-16 mice per group pooled from two independent experiments (unpaired Student’s *t* test). (**B-F**) In the Barnes maze, learning curves show mean daily distance run (**B**), and mean numbers of errors of holes visits (**C**) prior to finding the target hole connected to a hidden exit box. Effect of the time (p<0.0001) was analyzed by repeated measures mixed-effects models. **D**) Mean distribution of daily strategies used by mice to find the target hole in the course of the 6 training days and the probe trial are shown as percentages per group. Freeman-Halton extension of the Fisher’s exact test using 3 groups (mixed and spatial groups were merged) was used to compare distributions. **E**) Percent of time spent by mice in the target (T) quadrant *vs*. the three other quadrants (adjacent (A1 and A2), or opposite (Op)) during a 90-s probe trial run 24 h after the last training trial in the Barnes. Results were tested by the one-sample t-tests compared to chance (25%). **F**) Number of visits of quadrant containing the original exit hole (target) *vs*. other holes in other quadrants (other) during the probe trial. Results were analyzed by paired *t*-tests between target and other or by Student-t tests compared to the Saline group as indicated by brackets. *n* = 14–17 mice per group of treatment pooled from 2 independent cohorts. **G**) The novel object recognition task was performed 24 h after a training step with two identical objects. Bars represent the time spent by mice exploring the object that is novel (N) or not (Old, O), expressed as percentage out of the total time spent exploring either object. (**H-I**) The object location task was performed 3 h after the initial exploration on wildtype non transgenic mice (**H**, *n* = 13–14 mice per group of treatment pooled from 2 independent cohorts) and on *Il1r1*^neuronKO^ versus *Il1r1*^WT^ mice (**I**, *n* = 8–14 mice per group of treatment pooled from 2 independent cohorts) 24 days post-implantation. Each dot represents an individual value. Bars represent the time spent by mice exploring the object that was moved (M) or not (unmoved, U), expressed as percentages out of the total time spent exploring either object. Results were tested by one-sample t-tests compared to chance (50%). ns, non-significant, *p<0.05, **p<0.01, ***p<0.001. Graphs show means ± s.e.m.

To establish if the impact of IL-1 signaling on cognition is neuron-intrinsic and direct, we used the CaMKIIα-CRE-ERT2:*Il1r1*^flox/flox^ mice. We implanted IL-1β- or saline-infusing osmotic minipumps subcutaneously to *Il1r1*^WT^ and *Il1r1*^neuronKO^ mice and performed behavioral analyses. First, as in C57BL6/J mice upon chronic IL-1β infusion (Fig. 3F & H), we confirmed that IL-1β-infused *Il1r1*^WT^ mice were impaired in precision and consolidation of spatial memory in the Barnes maze and OL tests (Extended Data Fig. 2F and Fig. 3I). Remarkably, *Il1r1*^neuronKO^ mice infused with IL-1β performed as well as saline-treated mice, showing that knocking out *Il1r1* in glutamatergic neurons is sufficient to prevent the impact on memory, of chronic IL-1β administered systemically.

Together, these data indicate that chronic peripheral IL-1β impairs the spatial component of memory retrieval and consolidation, through direct signaling within excitatory glutamatergic neurons of the hippocampus.

### Chronic systemic exposure to low level of IL-1β induces leukocyte infiltration in the hippocampus without overt microglial activation

To assess how hippocampus-dependent cognitive impairments caused by chronic levels of IL-1β relate to changes in brain-resident or brain-infiltrated immune cells, we first analyzed the distribution and activation of non-neuronal cells such as astrocytes, microglia and peripheral immune cells by immunofluorescence microscopy and flow cytometry analyses. First, we confirmed that IL-1β is present in the hippocampus following chronic systemic infusion of IL-1β (Extended Data Fig. 4A). Chronic IL-1β seemed to only mildly impact microglia and astrocyte numbers (Extended Data Fig. 4B-D) and did not significantly impact microglia activation (Extended Data Fig 4E-G). However, chronic systemic low levels of IL-1β drove CNS infiltration of monocytes, neutrophils/granulocytes, and Ly6C^lo^ monocyte-derived cells, and T cells (Extended Data Fig. 4H-M), albeit with a lower magnitude than following *T. gondii* infection (see Extended Data Fig. 1). We also found that neither IL-1β nor the KO of neuronal *Il1r1* in *Il1r1*^neuronKO^ mice impacted on microglia or astrocyte numbers (Extended Data Fig. 5A-B). Since alterations in the neurogenic niche may underlie cognitive memory perturbations, we next assessed to which extent IL-1β affected the neurogenic niche. As an endpoint study, we counted the number of newly born doublecortin-positive neurons in the dentate gyrus of the hippocampus using immunohistochemistry analyses (Extended Data Fig. 5C). We found that neither chronic IL-1β, nor changes in neuronal IL-1R1 signaling altered the number of doublecortin-positive cells (Extended Data Fig. 5C), suggesting that the observed memory impairments of IL-1β-treated mice are unlikely to be due to changes in glial cell activation or newborn neurons.

### Chronic *T. gondii* infection and systemic exposure to IL-1β increase DNA double-strand break marker levels in the hippocampus

Because of the long-lasting impact of IL-1β on hippocampus function, we hypothesized that chronic IL-1β could act on neurons through an epigenetic mechanism. Thus, we chose to explore the role in this process, of the DNA DSB response, as a driver of epigenetic changes. As previously shown by us and colleagues, neuronal DSB levels can be reliably quantified by immunostainings of the phosphorylated form of the histone variant H2A.X (γH2A.X), or of 53BP1, two specific hallmarks of DSB in neurons, which form foci at the DSB sites ^40–42,55–57^. First, we counted the number of neurons from the dentate gyrus (DG) and the *Cornu Ammonis* (CA) bearing 53BP1-positive foci in *T. gondii*-infected mice (Fig. 4A). As shown previously ^55,58^, we confirmed using confocal immunofluorescence microscopy that mature neurons express diffuse 53BP1 staining in their nuclei (Fig. 4Ai , ii & Extended Data Fig. 4N). As part of the DSB response, 53BP1 form foci at sites of DSB, that can be identified as brighter immunofluorescent dots in the nuclei of neurons (Fig. 4Aii). We found that in the context of both latent *T. gondii* infection (Fig. 4Aiii,B) and encephalitis (Fig. 4C), there was a significantly increased number of neurons bearing 53BP1-positive foci in the DG of the hippocampus. We hypothesized that this effect may depend on chronic neuroinflammation caused by the persistent infection. Chronic IL-1β-treatment indeed elicited a similar increase in DSB-bearing neurons in the DG and CA of the hippocampi of the mice as in the context of *T. gondii* infection (Fig. 4D), suggesting that this cytokine may directly trigger DSB in neurons. To test this hypothesis, we established primary cultures of mouse postnatal hippocampal neurons, treated them with increasing concentrations of IL-1β for 5 hours and analyzed the levels of γH2A.X by Western blot. Fifty (50) ng/ml of IL-1β were sufficient to increase by over 40% the amount of γH2A.X in the culture, compared to vehicle-treated controls (Fig 4E). At this concentration, no sign of neuronal loss was observed.

**Figure 4:**
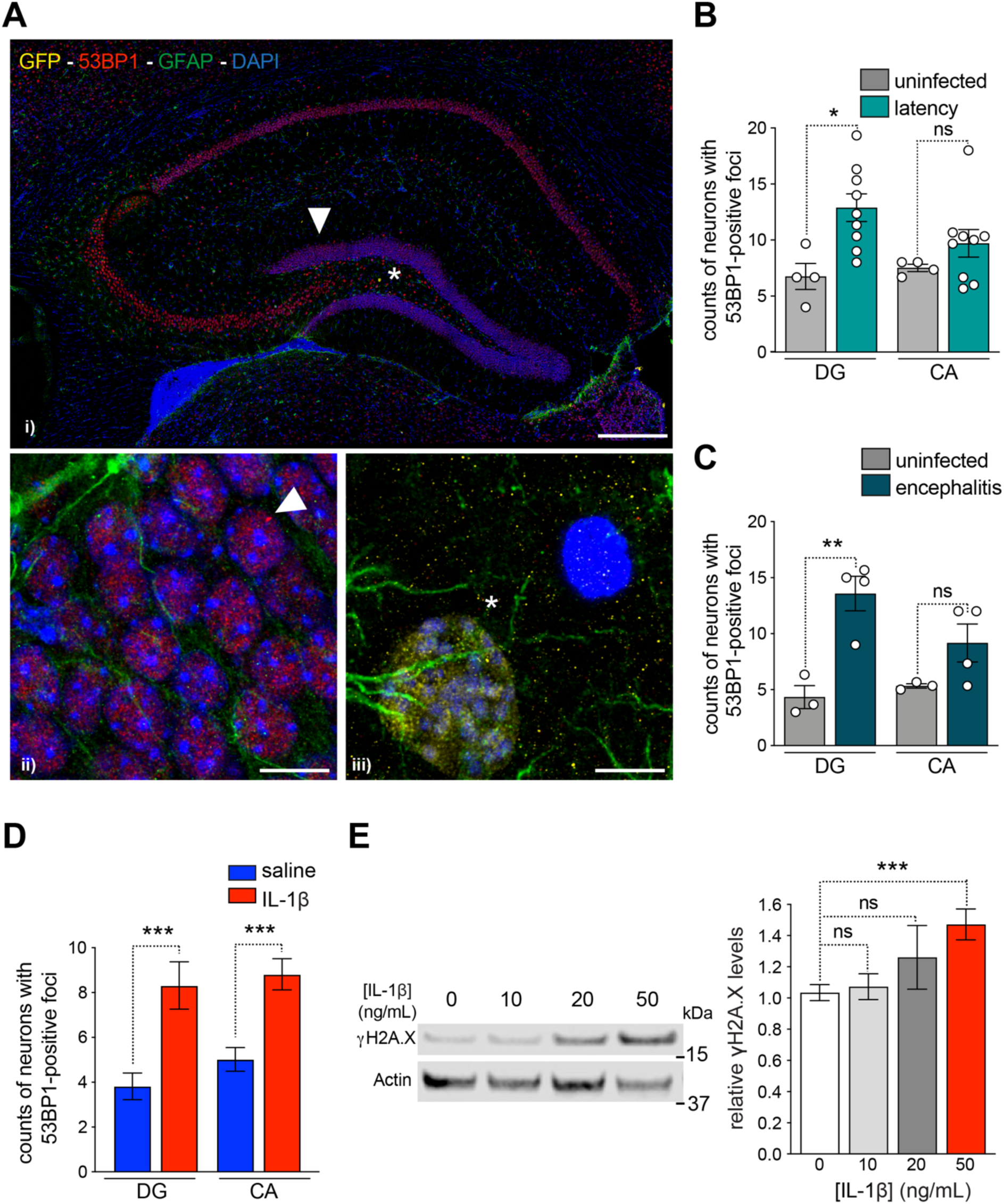
Chronic *Toxoplasma gondii* infection or exposure to IL-1β increase neuronal levels of DNA double-strand breaks. **A**) Confocal micrographs of the hippocampus of a mouse infected by latency-associated *Tg* strain (*Tg*.GRA6-OVA, in yellow), stained for DSB (53BP1, red), astrocytes (GFAP, green) and DAPI labeling of nuclei (blue). i) General view of the hippocampus indicating no changes in morphology and representative specific diffuse staining of 53BP1 in neurons. Scale bar, 200 μm. ii) Higher magnification view of granule cells in the DG showing a typical 53BP1-immunoreactive focus (red) in the nucleus of a neuron (white arrowhead in i)). iii) Detail of the rare presence of a *Tg* cyst in the hilus of the dentate gyrus (DG) (as shown by a star in i)). Scale bar, 20 μm. (**B-D**) DG cells and neurons in the CA1-3 with 53BP1-positive foci were counted and averaged on three coronal sections per mouse, upon chronic latent *Tg* infection (**B**, *Tg*.GRA6-OVA, *n* = 4–9 mice per condition), encephalitic *Tg* infection (**C**, Pru, *n* = 3–4 mice per condition) or at 28 days post-implantation of IL-1β-versus saline-infusing minipumps (**D**, *n* = 18-19 mice per condition). **E**) Cultures of primary hippocampal neurons from WT C57BL/6 mice were exposed to the indicated concentrations of mouse IL-1β or vehicle (0) for 5 h. Levels of the DSB marker γH2A.X were determined by Western blotting. The average γH2A.X signals to α-tubulin ratio in vehicle-treated cultures was defined as 1.0. *n* = 10–27 wells per condition from 11 independent experiments. In Western blots, each lane contained a sample from a different well of one culture. ns, non-significant, *p < 0.05, **p < 0.01, ***p < 0.001 *vs*. uninfected (**B, C**) or saline (**D**), or vehicle (**E**) by Kruskal-Wallis and Dunn’s multiple comparison post-hoc tests. Each dot represents the mean value for an individual (B-D) or a well (E). Bars represent means ± s.e.m.

Together, these data suggest that a DNA DSB response involving γH2A.X and 53BP1 is induced in hippocampal neurons of the DG upon *T. gondii* infection and upon chronic systemic exposure to IL-1β.

### H2A.X-dependent signaling critically contributes to memory deficits caused by *T. gondii* infection and by chronic exposure to IL-1β

We next tested whether the induction of a H2A.X-mediated DNA DSB response plays a role in the consolidation deficits of spatial memory in mice. Because the complete knockout of genes involved in the DSB response is often associated with embryo lethality or neurodevelopmental deficits caused by genome instability ^59–61^, we decided to target H2A.X expression selectively in neurons of adult animals. To this aim, we treated adult CaMKIIα-CRE-ERT2: *H2ax*^flox/flox^ mice with tamoxifen to invalidate the *H2ax* gene in excitatory neurons at adulthood. We verified that CaMKIIα-CRE-ERT2: *H2ax*^flox/flox^ treated with tamoxifen loose the expression of H2A.X in excitatory neurons (H2ax^neuronKO^ mice, Fig. 5A). Absence of H2A.X in excitatory neurons did not impact the general brain morphology, the global distribution of glial cells or the number of newborn neurons (Extended Data Fig. 5D-F). First, we implanted *H2ax*^neuronKO^ and control *H2ax*^WT^ mice with osmotic IL-1β *vs.* saline mini-pumps as previously, and tested the mice in the OL task (Fig. 5B). As expected, IL-1β impeded consolidation of spatial memory in *H2ax*^WT^ mice as indicated by the absence of preferred exploration of the moved object. Importantly, saline-treated *H2ax*^neuronKO^ mice had the same ability to consolidate memories as *H2ax*^WT^ mice exposed to saline (Fig. 5B), showing that under basal/non-inflammatory conditions, H2A.X expression in neurons is not required for the learning and the consolidation components of spatial memory. Notably, *H2ax*^neuronKO^ mice treated with chronic IL-1β were also able of memory consolidation, showing that knocking out H2A.X was protective against the deleterious impact of IL-1β on memory. Finally, we confirmed this finding in the more complex situation of *T. gondii* infection. In the OL task, we confirmed the *T. gondii*-induced crippling of consolidation processes in *H2ax*^WT^ mice (Fig. 5C,D). As above with IL-1β administration, knocking out *H2ax* in excitatory neurons was sufficient to prevent the deleterious impact of *T. gondii* encephalitis on consolidation of spatial memory (Fig. 5C), without affecting the brain parasite burden (Fig. 5D).

**Figure 5:**
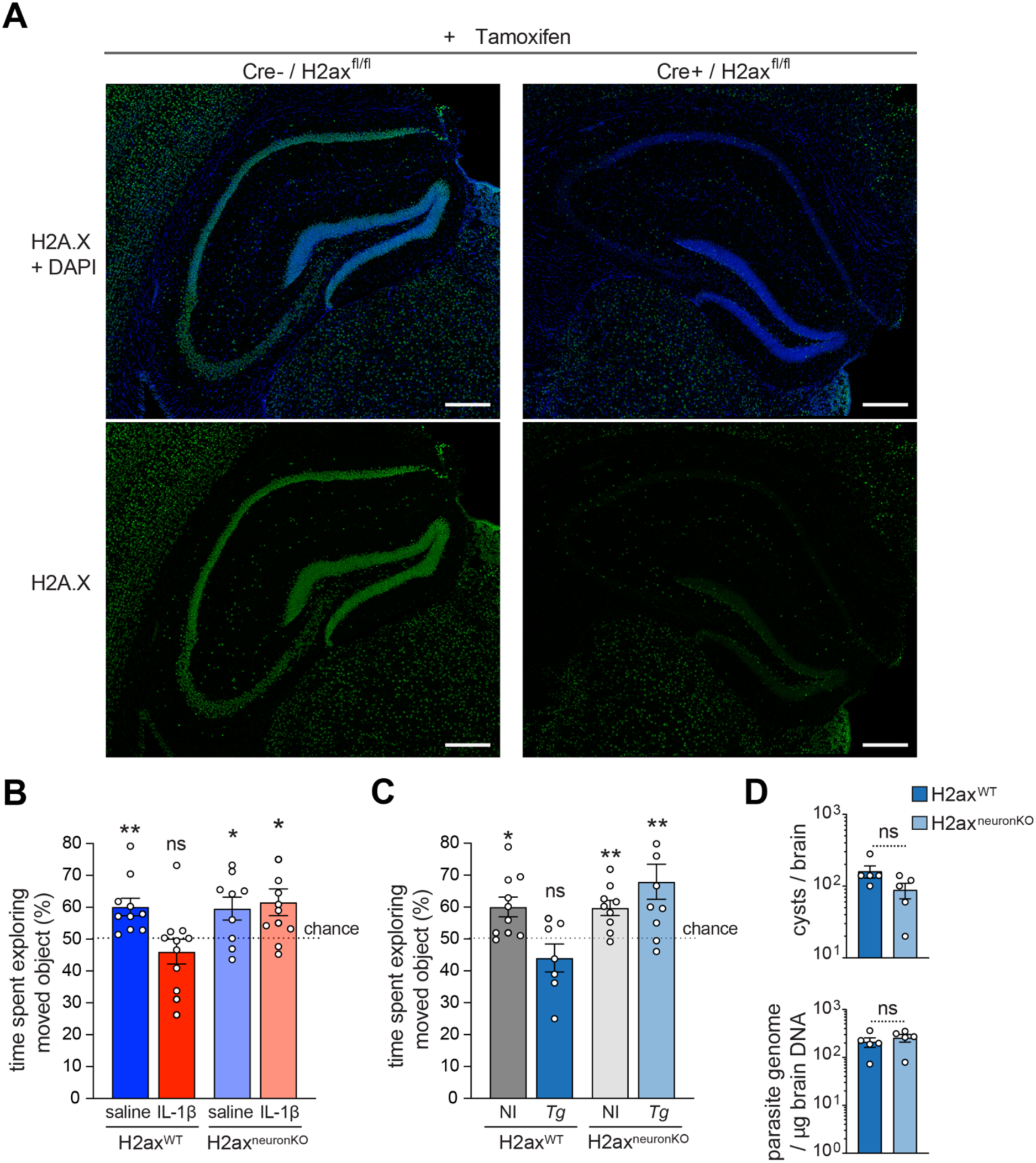
Knocking out *H2ax* in excitatory neurons is sufficient to prevent the deleterious effects of chronic *Toxoplasma gondii* infection and exposure to IL-1β on memory consolidation. A) Representative micrographs showing the total H2A.X (in green) and nuclear DAPI (blue) signals in the hippocampi from one *H2ax*^WT^ (*Cre^-^:H2ax^fl/fl^*) and one *H2ax*^neuronKO^ (*Cre^+^:H2ax^fl/fl^*) mouse. Both mice received tamoxifen, belonged to the same cohort and were implanted with saline-infusing minipumps. Scale bar, 200 μm. (**B-C**) The object location task was performed 3 h after an initial exploration of the objects with *H2ax*^WT^ and *H2ax*^neuronKO^, 10 weeks after tamoxifen treatment, and either 24 days post-implantation of IL-1β-versus saline-infusing minipumps (**B**, *n* = 9-11 mice per condition from 2 independent cohorts) or 5 weeks post-infection by encephalitic *Tg* strain ME49 (*Tg*) *vs*. uninfected (NI) controls (**C**, *n* = 7-10 mice per condition pooled from 2 independent cohorts). **D)** Cyst number and total parasite burden in the brain was assessed in one experiment (*n* = 5 mice per condition). Bars represent the time spent by mice exploring the object that was moved, expressed as percentage out of the total time spent exploring either of the two objects. Results were tested by one-sample t-tests compared to chance (50%) (B, C) or by Student’s t-tests (D). ns, non-significant, *p<0.05, **p<0.01. Each dot represents an individual value. Values are means ± s.e.m.

Taken together, our results show that blockade of the H2A.X-mediated DNA DSB response is protective against the IL-1-induced impairment of spatial memory consolidation not only in a simplified model of IL-1β systemic infusion but also in the relevant and prevalent context of CNS infection by *T. gondii*.

## DISCUSSION

Our study shows that IL-1 signaling in excitatory neurons contributes to memory impairment in two chronic pathological contexts leading both to an increase in IL-1 in the hippocampus (i.e. a brain-persisting infection by *T. gondii* parasites, and a chronic exposure to systemic IL-1β). Additionally, we uncover that DNA DSB signaling mediated by H2A.X in neurons is a critical mechanism involved in the IL-1-induced spatial memory deficits. We indeed found that the IL-1-induced memory impairment could be completely prevented by the invalidation of H2A.X, a histone variant involved in the DNA DSB response. This provides an unprecedented proof of concept that the IL-1 and H2A.X signaling pathways may be relevant therapeutic targets to mitigate memory dysfunction in neuroinflammatory pathologies. A major asset of this work is to identify a novel mechanism that links cytokine (IL-1)-regulated neuronal epigenetic processes and chronic inflammation-induced memory disorders.

Several questions arise, including the type of signaling events between IL-1R1 & DSB, and the nature of the H2A.X-mediated processes driving hippocampal neuron dysfunction & spatial memory disturbances.

Regarding the signaling pathway driving DSB formation and recognition downstream of IL-1R, previous data have suggested that pro-inflammatory cytokines can alter neurotransmission ^22,25,62,63^ and that disturbed neurotransmission contributes to DSB accumulation by shifting the repair/breakage equilibrium ^41,42^. Therefore, it is most likely that neurons exposed to IL-1 may display disturbance of neurotransmission resulting in DSB accumulation. Strengthening this hypothesis is the well-known close functional interaction of IL-1R1 with the ion-channel receptor to the glutamate neurotransmitter, the N-methyl-D-Aspartate receptor (NMDAR) in the hippocampus ^63^. Indeed, acute exposure to cytokines including IL-1β, results in impaired reuptake of glutamate. Notably, the uptake of glutamate was also found to be dysregulated upon chronic *T. gondii* infection ^64^. This, in turn, may lead to high levels of glutamate, causing abnormal activation of NMDARs excluded from the synaptic cleft (also designated as extrasynaptic NMDARs such as, in particular, GluN2B subunit-containing NMDAR, ultimately resulting in excitotoxicity ^25,62,63,65^. Next, IL-1R1 and GluN2B subunits of NMDAR co-localize in the postsynaptic density of dentate granule cells in the rat hippocampus, and IL-1β stabilizes this subunit at the cell surface, which is thought to contribute to increase the pool of extrasynaptic NMDAR^66^. Also, co-stimulation by IL-1β of neurons activated by NMDA facilitates NMDA-induced cation [Ca^2+^] increase and NMDAR-dependent excitotoxicity^63^. Moreover, in mouse neurons, the activation of extrasynaptic GluN2B-containing NMDAR leads to DSB accumulation by causing an imbalance in the DSB formation/repair process through promoting the fast degradation of DNA DSB repair factor BRCA1 ^40–42^. Hence, this mechanism may be at play in the findings we report here.

At this stage, it is unknown whether this phenomenon is specific to the IL-1 pathway or whether other cytokine signaling pathways also induce an excess of DSB accumulation and/or DSB signaling. To address this question, it would be useful to analyze how neurons respond to cytokines which signal through pathways that are either partially overlapping with the IL-1 pathway (e.g. IL-18 or IL-33), or that are largely different from it (e.g. IFN-I, IFN-γ).

The fact that knocking out neuronal *H2ax* is sufficient to prevent chronic spatial memory dysfunction in our two models is another original finding of our work. The mechanisms remain elusive but we envisage two non-mutually exclusive working models. In a first model, the neurons are both the receivers of the cytokine signals and also the sole cells responsible for memory dysfunction. Work by us and colleagues has previously demonstrated that a balance between DNA DSB formation and repair is critical in cognitive processes, and that dysregulations leading to accumulation of DSB in neurons impair memory formation and participate to age-related neurodegenerative processes ^40–42,67,68^. We hypothesize that by knocking out *H2ax* in excitatory neurons, γH2A.X foci were prevented to build up around DSB in neurons and the pathological consequences of their persistence were blocked. Under homeostatic circumstances, physiological activity-dependent neuronal DSB are constantly produced and repaired rapidly after generation. This process was shown to affect preferential regions of the neuronal chromatin, and was mostly considered as changing the expression of memory genes ^40–42,56,57,67–69^. At this stage, it is still not understood if, under inflammatory conditions, the same chromatin sequences are altered when unrepaired DSB accumulate and/or γH2A.X foci persist.

Our second working model comes from the observation that chronic systemic exposure to IL-1β is sufficient for leukocyte (including monocytes and T cells) recruitment in the hippocampus, and supports the idea that other (neuro)immune cells may also contribute to spatial memory impairment. These findings suggest an alternative model in which IL-1, by activating H2A.X-dependent signaling in neurons, may alter the cross-talk of excitatory neurons with surrounding neuroimmune cells and, fuel a vicious circle of deleterious changes in neuronal gene expression and function, through the action of these neighboring cells. This would be in line with recent findings that γH2A.X foci-bearing neurons promote neurodegeneration-associated profiles in neighboring microglia ^70^ and would be reminiscent of a previous study in a murine model of acute encephalitis showing that neurons responding to IFN-γ produce chemokines to attract phagocytes, including activated microglia, which in turn impede neurotransmission resulting in motor impairment ^71^. In this alternative working model, neurons are also the main receivers of IL-1 signals but additional neuroimmune cells beyond neurons would play a deleterious role in spatial memory.

Regardless of the model, neuronal H2A.X-related signaling remains key to the cognitive performance. Alike defective DNA repair, which cause cognitive deficits, the absence of breakage of the DNA during learning processes also impairs memory formation ^40,56,67,68^. For example, decreasing Topoisomerase 2β levels, an endonuclease which induces DSB in neurons upon activity, decreases DSB levels together with γH2A.X levels and compromises fear learning ^40,72^. Yet, until now, manipulations of neuronal DSB response were not able to dissociate the effects of the breakage of the DNA itself from effects of its associated epigenetic signaling. Here, we provide the first demonstration that neuronal H2A.X, as part of DSB epigenetic signaling, is dispensable for the physiological consolidation of spatial memory, yet is detrimental when its accumulation lasts in pathological contexts. Future investigations should help define the ultimate events mediated by H2A.X that are imbalanced in spatial memory disturbances.

*T. gondii* infection notoriously changes innate behaviors, notably by enhancing risk-taking behavior ^20,73,74^, which involves other brain areas such as the amygdala. To which extent the IL-1/H2A.X-dependent mechanism described here could explain enhanced risk-taking behavior linked to *T. gondii* infection, remains to be investigated. Due to constitutive IL-1R1 expression uniquely restricted to the dentate gyrus ^48^ (https://mouse.brain-map.org/gene/show/15950)^75^, one could suspect that the dentate gyrus be more sensitive to IL-1 than other brain areas. Nonetheless, the hippocampus is not a structure involved in risk assessment behaviors. Conversely, signaling by other pro-inflammatory cytokines, such as IFNs or chemokines, which are produced massively in the brain during *T. gondii* infection, may be more prominently involved in the dysfunction of other brain regions ^24^. Altogether, it is tempting to speculate that the nature of the cytokines involved in *T. gondii*-induced neurological dysfunction will vary according to the brain area implicated and the type of immune receptors expressed or induced in neurons of the concerned area. In any case, it is fascinating that although *T. gondii* infection elicits a large array of neuroinflammatory cells and pathways in the brain, its effects on spatial memory consolidation could be pinpointed to the single IL-1 pathway.

Regardless of the final mechanisms, our data clearly support a model whereby *T. gondii*-induced cognitive dysfunction is caused chiefly by neuroinflammation and is not the result of a direct action of *T. gondii* on infected neurons. Yet, our data do not rule the possibility that the capacity of *T. gondii* to modulate host cytokine signaling ^76^ might also influence the cerebral response and, ultimately, affect the magnitude of some cognitive changes. Interestingly, GRA15, a parasite protein that increases NF-kB signaling ^77^, has been shown to increase brain IL-1β RNA levels, and to sensitize mice to chemically induced IL-1-dependent seizures upon *T. gondii* encephalitis ^78^. Further studying the role of GRA15 or TEEGR, a parasite-secreted effector that, conversely, negatively modulates IL-1/NF-kB signaling ^79^ in the context of latent *T. gondii*, could be useful to better understand the role of parasite factors in spatial memory dysfunction.

Up to now, the impact of IL-1 on cognitive function has been mostly investigated after acute exposure to the cytokine, which is known to induce sickness behavior syndrome (SBS). In severe *T. gondii* infection, IL-1 receptor signaling was shown to promote chronic cachexia ^49^. Signaling through endothelial IL-1 receptor was reported to be the key driver of SBS, and at the cellular level, of microgliosis and impaired neurogenesis ^48,80,81^. So far, behavioral studies of chronic IL-1β exposure only focused on sleep regulation ^82^, aberrant mood control and activity ^25,53^, and depressive innate behaviors^53^. Remarkably, in our hands, the low concentration of IL-1β we used did not lead to SBS, as assessed in the EPM, in the habituation components of the object location test, or in the general exploratory activity and distances in the Barnes maze. At the cellular level, chronic IL-1β did not have a major effect on neurogenesis ^53^, and it only slightly increased microglial numbers with no change in the activation markers that we evaluated. Therefore, our study pioneers the breakthrough idea of a SBS-independent, specific and direct impact of chronic low levels of IL-1β on spatial memory consolidation and precision memory, two components specifically encoded by the dentate gyrus of the hippocampus. Altogether, this suggests that effects of the chronic presence of cytokines involve a ballet of cellular signaling pathways in brain structures, that differ from the acute situation and is choreographed depending on the local concentration of the cytokine.

Importantly, beyond infectious contexts, chronic mild elevated levels of plasmatic IL-1β have been reported in several chronic neuropsychiatric diseases, such as depression, bipolar disorder, or schizophrenia^83–86^. Interestingly, a role for latent *T. gondii* infection in these pathologies has been suggested ^6–10^, and the impact of *T. gondii* on cognitive processes may have been easily overlooked given that our findings in mice demonstrate fine defects in the consolidation and precision components of spatial memory. Nonetheless, this impact on precision may be critical to navigation and truly disabling in everyday tasks if confirmed in humans. In addition, elevated IL-1β concentration has been detected in the plasma and the brain both of patients suffering from Alzheimer’s disease ^87^ and in animal models of AD ^88,89^. Therefore, our results represent a potential breakthrough for the understanding of the role of IL-1 in memory deficits occurring in these pathologies. Our work could have major ramifications in a myriad of neurological disorders where IL-1 is upregulated, either systemically or locally in the CNS. Of note, it was recently shown that DSB signaling in neurons drastically reprogram neuroinflammatory microglia toward neurodegeneration-associated phenotypes ^70^. Exploring the therapeutic potential of manipulating the neuronal DNA DSB response could thus provide additional therapeutic prospects to alleviate inflammaging-related diseases.

## MATERIAL AND METHODS

### Mice, infection and tissues sampling

Specific pathogen-free C57BL/6J male mice were purchased from Janvier Laboratories (France) at 6-8 weeks of age. Mice were acclimated for 1-2 weeks in the experimental BSL2 facility. We also studied CaMKIIα-CRE-ERT2^+/-^: Il-1r1^flox/flox^ and CaMKIIα-CRE-ERT2^+/-^: H2ax^flox/flox^ transgenic male mice. CaMKIIα-CRE-ERT2^-/-^ (Cre negative) littermates of each strain were use as controls. All mice were on a pure C57BL6/J background (see refs. ^1–3^). Transgenic mice were bred at the UMS006-ANEXPLO-CREFRE facility (Toulouse). To induce the knockout of the floxed gene, 6-8 week-old transgenic mice were injected daily intraperitoneally (ip) by a suspension of tamoxifen (Sigma-Aldrich) in corn oil (Sigma-Aldrich) for 5 days at 10 mg kg^-1^ day^-1^. Commercial and transgenic mice used in this study were implanted with osmotic minipumps (Alzet AZT2004, Charles River) at 8 weeks of age or were injected intraperitoneally (i.p.) with 200 tachyzoites or mock-injected with PBS (uninfected group) between 9 and 16-weeks of age. Behavior assessments were carried out starting 24 days post-implantation or between 6 and 16 weeks post-infection.

Mice were acclimated for one week in the experimental ABSL2 facility, prior to intraperitoneal (i.p.) injection with 200 Pru- or 2000 ME49-derived tachyzoites in 200 µL of PBS, or with 200 µL of PBS only for the uninfected group. During acute infection, mice were observed daily and dietary supplement (DietGel® Recovery, Clear H_2_O) was added in all experimental conditions. Mice were routinely weighed throughout infection.

Littermates were group-housed. For testing in the Barnes maze, mice were single-housed from 3 days before the beginning of training until the test was concluded. All mice were housed in cages with enrichment (nesting material and shelter) at 24°C, had *ad libitum* access to irradiated A4 food (R04-10, SAFE Diets) and water and were exposed to a 12-h light/dark cycle. For histological and biochemical analyses, mice were anesthetized with Avertin (tribromoethanol, 250 mg kg^-1^, ip) for IL-1β infusion experiments or with a mix of ketamine (200 mg kg^-1^) and xylazine (20 mg kg^-1^) injected i.p for parasite-related experiments and assessments of the immune response. Mice were then perfused transcardially with 0.9% NaCl. One hemibrain was used fresh for parasite load analysis or flow cytometry, or snap frozen and stored at –80°C for bulk RNA sequencing, cytokine immunoassay or biobanking. The other hemibrain was drop-fixed in 4% paraformaldehyde in phosphate-buffered saline (PBS) and sectioned (30 μm) with a sliding microtome (Leica SM2000R). All mouse experiments were performed in accordance with European Union Council Directive 86/609/EEC, and experiments were performed following the French national chart for ethics of animal experiments (articles R 214-87 to -90 of the “Code rural”). Our protocols received approvals from the local committee on the ethics of animal (CEEA14) and the AFiS cell of French Ministry of Research.

### *Tg* parasite strains and culture

Derivatives of type II *Tg* strains (Prugniaud and ME49) were used in this study (see Table 1 for details). For both strains*, Tg* tachyzoites were maintained *in vitro* by serial passage on confluent monolayer of human foreskin fibroblasts (HFF, ATCC SCRC-1041) in Dulbecco’s Modified Eagle Medium (DMEM, Gibco) supplemented with 1% (vol/vol) fetal bovine serum (FBS, Gibco), 1% (vol/vol) penicillin/streptomycin (Gibco), 0,1% (vol/vol) 2-mercaptoethanol (Gibco). For *in vivo* infections, parasites were prepared immediately before mouse infection. Infected HFF were scraped and remaining intracellular tachyzoites were released through homogenization with a 23G needle. Freshly egressed parasites were filtered through a 3 µm polycarbonate hydrophilic filter (it4ip S.A.), centrifuged at 1000 g for 10 min and resuspended in phosphate-buffered saline (PBS, Sigma-Aldrich). Tachyzoites were individualized through a 25G needle, counted and diluted to 10^3^ or 10^4^ parasites/ml.

**Table 1:** List of transgenic Prugniaud and ME49 derivative strains used in this study.

| Short name | Complete description, with full genotype | Reference |
| --- | --- | --- |
| Pru | Pru.Δhxgprt.tdTOMATO<sup>prom TUB</sup> | 4 |
| Tg.SAG1-OVA | Pru.Δhxgprt.tdTOMATO<sup>prom TUB</sup>.SAG1ΔGPI-OVA<sub>[140-386]</sub><sup>prom TUB/3'utr DHFR</sup> +BLE | 4 |
| ME49 | ME49.Δhxgprt.RFP | 5 |
| Tg.GRA6-OVA | Pru.Δhxgprt.GFP<sup>prom GRA1</sup>.click beetle LUC<sup>prom DHFR</sup>.GRA6<sub>II</sub>-LEQLE-SIINFEKL<sup>prom GRA6/3'utr</sup> GRA2 +HXGPRT | 6 |

### Osmotic minipump implantation

Mouse recombinant IL-1β (Immunotools GmbH, Germany) was dissolved in sterile saline solution (0.9% NaCl) containing 0.1% Bovine serum albumin as a carrier. For the chronic experiments, mice were implanted subcutaneously (s.c); in the interscapular region with osmotic minipumps (model AZT2004, Alzet, Charles River). Minipumps were filled with saline (saline + carrier) or IL-1β solution per the manufacturer’s instructions. Model 2004 minipumps delivered fluid at a rate of 0.25 μL/h for up to 35 days. Concentration of IL-1β in each minipump was calculated to infuse 5 μg. kg^-1^. day^-1^ ^7^. After 24h of priming in saline solution at 37°C, minipumps were surgically implanted in mice anesthetized with Avertin (tribromoethanol, 250 mg. kg^-1^, AlphaAesar, Merck). S.c. injections of 0.05 mg. kg^-1^ of Buprenorphine (AnimalCare) were use perioperatively to prevent pain during the recovery of the mice. Two (2) weeks post-implantation, to prevent minipump expulsion, mice were lightly anesthetized with 4% Isoflurane, potential skin adherences at the tip of the pumps were disconnected and a surgical clip (Reflex clip 9 mm, WPI) was placed instead.

### Behavioral testing

For all behavioral tests and subsequent analyses, experimenters were blinded to the genotype and treatment of mice. Mice were assessed in the following tests in the indicated sequence and as described in the corresponding references: elevated plus maze (EPM) ^8^, object location (OL) and novel object recognition ^8,9^ and the Barnes maze, with the following modifications. In the EPM, videotracking and analysis were performed using ANYMaze software (v7.08, Ugobasile, Italy). In the OL and the NOR, after an initial step of exploration of two identical objects in a large arena (60 x 30 cm) for 10 minutes, mice were allowed to explore again the arena 3 hours later for 10 minutes, when one of the familiar object had been moved (OL). Twenty-four (24) hours later, one familiar object was replaced by a novel object in terms of shape, texture and colors, and mice were allowed to explore the arena for 10 minutes. Activity of the mice was videorecorded using camcorder and videos converted into mp4 format and subsequently analyzed: i) automatically by a homemade software (coded with Python) to exclude any side preference bias and to record the total distance ran ; ii) manually analyzed using a free video annotation tool (ANVIL 6, http://www.anvil-software.de/, Germany) to determine times and pokes of exploration of the objects. More precisely, a mouse was considered as exploring an object when its nose was in contact with the object and its four paws were on the ground. In the Barnes maze, videotracking and analysis were performed using Ethovision XT15 software (Noldus, USA). A cued-box trial preceded the hidden-exit box trainings, consisting in a first trial of releasing the mouse in the middle of the maze for 3 minutes, with an exit box linked to the target hole, hidden under the table of the maze with a reward cereal loop inside it. Passed the three minutes of each trial, if the mouse had not found the exit box, it was guided by the experimenter to it and received a cereal loop as a reward as soon as it had found the box. Four hidden-exit box trainings trials per day for *Tg*-related experiments and 3 trials per day for pumps-related experiments were performed during 5 days to allow each control groups to reach a steady plateau in their performances without overtraining. Cohorts of mice implanted with osmotic minipumps underwent either EPM, OL and NOR or EPM and Barnes maze.

### Immunocytochemistry

Immunostaining of mouse brain sections was performed essentially as described ^8^. TBS was used and steps were added for antigen retrieval and peroxidase inhibition to enhance nuclear staining. Blocking solution was 5% normal goat serum in 0.1% TBS-Tween20. Sections were incubated overnight at 4 °C with primary antibodies, including monoclonal mouse anti-GFAP (clone GA5, Millipore) or polyclonal rabbit anti-GFAP (Dako); polyclonal chicken anti-GFP and rabbit anti-53BP1 (Novus Biologicals), rabbit anti-doublecortin (Synaptic systems), rabbit anti-IBA-1 (Wako) and chicken anti-MAP2 (ABCam). Primary antibodies were diluted 1:500 (or 1:1000 for anti-doublecortin) in blocking solution. Secondary antibodies were goat anti-mouse or anti-rabbit Alexa 488, anti-rabbit Alexa 546, anti-rabbit Alexa 633 or Alexa 647 (ThermoFisher). Biotinylated anti-rabbit antibodies (Vector Laboratories) were used for doublecortin. Diaminobenzidine (Vector Laboratories DAB kit) was used as a chromogen. After extensive rinses, coverslips or sections were mounted with Prolong gold mounting medium (ThermoFisher) or with Fluoromount medium (Electron Microscopy Sciences).

For quantification of GFAP-positive or IBA-1-positive signals on DAPI, digitized images of dentate gyrus region were obtained with a digital brightfield Olympus BX41 microscope with a DP11 digital camera system and a X-cite120 Q (Lumen Dynamics) lamp or with Zen blue v3.1.0.00007 software (Zeiss) on a Zeiss Axio-observer widefield microscope. Image were manually annotated with cell counter tools from Image J, over three frames per mouse. Counts were converted into densities using the area covered by the counting in mm^3^.

Tiles of z-stacks of images of 53BP1 foci or GFP and IBA-1 and GFAP signals were obtained with a Leica SP8 confocal microscope, and a 20X objective. Images were acquired and processed with LSM LAS X 4.0.2 software (Leica). Z-stacks of merged mosaic confocal images were processed with Image J, and final images were obtained by maximal-intensity projection.

53BP1, and doublecortin stainings in the DG from coronal sections of mice were quantified essentially as described^10^. Counts of neurons positive with 53BP1 foci were averaged from 3 sections per mouse unless stated otherwise. Counts of doublecortin-positive neurons were averaged from 2-3 sections per mouse.

### Cell cultures and treatments

Primary cultures of hippocampal neurons were established as described ^9,10^, using postnatal day 0 pups from C57BL6/J WT mice. Neurons were plated on 12-well plates (Corning) at 0.5 x 10^6^ cells/well or on 12-mm glass coverslips coated with poly-D lysine (Millipore). Cultures were used for experiments at 14 DIV. Experimenters were blinded to treatments. Recombinant IL-1β (Immunotools GmbH, Germany) was dissolved at a stock concentration of 100μg/ml in sterile Phosphate Buffer Saline containing 0.1% Bovine serum albumin as a carrier. Various concentrations were applied to neurons for 5 hours as described^10^.

### Western blot analysis

Cultured cells were washed in PBS and scraped into RIPA buffer according to^11^. Briefly, lysates were sonicated for 10 min at 4°C and spun in a refrigerated microcentrifuge at maximal speed for 10 min. Protein concentrations were determined by Bradford assay. Proteins (20 μg per mouse sample) were loaded on 4–12% Bis-Tris gels (ThermoFisher), separated by SDS-PAGE, and transferred to nitrocellulose membranes. After 1 h in blocking solution (5% nonfat milk in Tris-buffered saline (TBS)), membranes were incubated with primary antibodies in 5% nonfat milk in TBS/0.1% Tween for 3 h at room temperature for mouse phospho-Histone H2A.X (ser139) monoclonal antibody, dilution 1/1,000 (ThermoFisher, MA1-2022) and mouse anti-α-Tubulin monoclonal, dilution 1/10,000 (Sigma, T6199) ; Blots were washed 3 times in TBS/0.1% Tween and incubated with secondary IRD-tagged antibodies (CF770, Biotium) diluted 1:10,000 in Odyssey blocking buffer for 1 h at room temperature.

Western blot signals were analyzed using an Odyssey Li-COR infrared imaging system (v1.0.37) coupled with Image studio 5.2 software (Li-COR). All protein signals were normalized on α-Tubulin signal.

### Hippocampus isolation and cell preparation for flow cytometry analyses

Hippocampus regions of the brain were microdissected and collected in cold Hanks’ Balanced Salt Solution (HBSS, Sigma-Aldrich), on ice-cooled plate under dissecting binocular. Hippocampus was then sliced with a scalpel and digested for 1 h at room temperature in HBSS supplemented with 0.5 mg/ml collagenase D (Roche) and 10 µg/ml DNase I (Sigma-Aldrich). During enzymatic digestion, tissue was mechanically dissociated by trituration with plastic pipette every ten minutes. Collagenase activity was stopped by adding 10% (vol/vol) FBS in digestion mix, then samples were filtered through a 70 µm strainer (Falcon), washed with HBSS supplemented with 5% (vol/vol) FBS, 1% (vol/vol) Hepes (Gibco) and 1% (vol/vol) penicillin/streptomycin (complete HBSS or cHBSS). Samples were centrifuged at 600 g for 5 min, the cell pellet was resuspended in 30% (vol/vol) Percoll (Cytiva) and centrifuged at 1590 g for 30 min. Pellet was washed in cHBSS and centrifuged at 470 g for 5 min. Finally, remaining erythrocytes were lysed using ACK buffer (100 µM EDTA, 160 mM NH_4_Cl, 10 mM NaHCO_3_).

To saturate Fc receptors and detect dead cells, samples were incubated respectively with FcR Block (Biolegend) and eFluor780 Fixable Viability dye (1/1000, Invitrogen) diluted in PBS for 15 min at 4°C. Then, cell suspensions were surface-labelled with the following antibodies : CD45 PerCP-Cy5.5 (30-F11, 1/300, BD Pharmingen), CD11b PE-CF594 (M1/70, 1/3000, BD Horizon), CD3ε BV421 (145-2C11, 1/300, BD Horizon), Ly6G BV510 (1A8, 1/200, Biolegend), Ly6C BV711 (HK1.4, 1/1800, Biolegend), MHC II I-A/I-E FITC (2G9, 1/300, BD Pharmingen), CD86 APC (GL1, 1/300, BD Pharmingen) in PBS supplemented with 0,5% (vol/vol) FBS and 0,4% (vol/vol) EDTA (Sigma-Aldrich) for 30 min at 4°C. Samples were fixed for 20 min at 4°C in 4% paraformaldehyde solution (Electron Microscopy Sciences) diluted in PBS, acquired on a BD Fortessa X20 cytometer and analyzed using FlowJo (TreeStar, v10.7.1).

### Bulk RNA sequencing

Microdissected hippocampi were disrupted with ultra-turrax® in ice-cold QIAzol Lysis Reagent (Qiagen). Total RNA was purified with RNeasy Lipid Tissue Mini Kit (Qiagen). Libraries were generated with the Illumina® Stranded Total RNA Prep, Ligation with Ribo-Zero Plus kit and the IDT ILMN RNA UDI B Lig 96 Idx 96 Spl index kit (Illumina) following the manufacturer’s instructions. Briefly, RNA quantity was normalized, ribosomal RNA was depleted then RNA was fragmented and denatured. After cDNA synthesis, anchors were ligated and libraries were amplified.

For differentially expressed gene analyses, raw sequencing reads were processed using nf-core/rnaseq pipeline (v3.2) ^12,13^. Briefly, quality of raw sequencing files was checked using fastqc (v.0.11.9) [https://www.bioinformatics.babraham.ac.uk/projects/fastqc/], reads were trimmed using trimgalore (v.0.6.6) [https://www.bioinformatics.babraham.ac.uk/projects/trim_galore/] to remove Illumina universal adapter sequences. Quality of trimmed reads was assessed using fastqc (v.0.11.9) and indicated complete removal of adapter sequences. Trimmed reads were aligned to GRCm39 mouse genome reference assembly and genome annotation from Ensembl (mm39, release 103) using STAR (v.2.6.1d)^14^ and read quantification was performed using salmon (v.1.4.0) ^15^. Exploration and differential expression analysis of RNAseq data was performed using R [https://www.R-project.org/] (v.4.1.3) / RStudio (v.202202.0, build 443) and DESeq2 R package (v1.34.0) ^16^ using a custom script. Salmon transcript-abundance estimates were imported using the tximport R package (v 1.22.0) ^17^ and summarized at the gene level using the ‘salmon_tx2gene’ file generated by nfcore/rnaseq pipeline. Raw count matrix was filtered to remove lowly expressed genes. Genes having a minimum of 10 normalized reads in 3 of the 4 replicates in any of the Uninfected, Latency or Encephalitis samples were kept. Principal component analyses and hierarchical clustering using variance-stabilized normalized counts indicated that samples clustered together according to their group. Differential expression analyses were performed using DESq2::DESeq() function and extracted using DESq2::results() function with a ‘lfcThreshold’ parameter of 1 and an ‘alpha’ parameter of 0.05 (i.e. adjusted p-value ≤ 0.05). Functional enrichment analyses were performed using clusterProfiler (v 4.2.2) ^18^ R packages.

### IL-1β quantification

Serum IL-1β levels were assessed by ELISA according to the manufacturer’s instructions (Quantikine ELISA kit, R&D Systems). Microdissected hippocampi were homogenized with the FastPrep-24™ Classic Instrument in Lysing MatrixD tubes (MP Biomedicals) containing ice-cold cell lysis buffer (Invitrogen) supplemented with protease inhibitors (Thermo Fisher Scientific) and 1mM of PhenylMethylSulfonylFluoride (PMSF, Sigma-Aldrich) solution prepared in 2-Propanol (Sigma-Aldrich). Tissue lysates were centrifuged at 15 800 g for 15 min at 4°C, and cleared supernatants were collected and stored at -80°C until IL-1β immunoassay. Total protein concentration of the cleared lysate was assessed with the Pierce™ BCA Protein Assay Kit (Thermo Fisher Scientific). IL-1β was quantified from hippocampi tissue lysates with V-PLEX Mouse IL-1β Kit (Meso Scale Discovery) following the manufacturer’s instructions.

#### Whole brain parasite load analysis

Hemibrains were homogenized in PBS using Potter tissue grinder, then 10% of the homogenate was used either to assess total parasite load (tachyzoite and bradyzoite parasite stages) by qPCR or to enumerate cysts by visual counting as described earlier ^6^. Cyst wall was labelled with rhodamine or fluorescein-conjugated *Dolichos Biflorus Agglutinin* (DBA, Vector Laboratories) and cysts were counted using an inverted fluorescence microscope at X20 magnification. For total parasite load quantification, genomic DNA was extracted using the DNeasy Blood & Tissue Kit (Qiagen) according to the manufacturer’s instructions and a 529-bp repeat element in the *T. gondii* genome was amplified using the TOX9 (5′-AGGAGAGATATCAGGACTGTAG) and TOX11 (5′-AGGAGAGATATCAGGACTGTAG) primers ^19^. The number of parasite genome per μg of brain DNA was estimated by comparison with a standard curve, established with a known number of Prugniaud (Pru) tachyzoites.

### Blind-coding

Investigators who obtained data were blinded to infection and treatment of cell cultures and to genotype and treatment of mice. Investigators who analyzed the data were double-blinded.

### Statistical analysis

GraphPad Prism software (v8.4.0) and R studio (v4.1.0) were used for statistical analysis. Data normal distribution was determined with Agostino & Pearson normality test, and the Bartlett-test was used to verify equality of variance. Welch’s correction was applied when the variances were not equal. Parametric or non-parametric tests and corrections for multiple comparisons were described in the legend of each figure. To assess whether differences observed in strategy types used during the Barnes maze were statistically significant between infected and uninfected groups, a Freeman-Halton extension of the Fisher’s exact test was performed on strategies (mixed and spatial groups were fused to allow appropriate use of the test since mixed strategy is partly spatially-driven). Null hypotheses were rejected at the 0.05 level.

## DATA AVAILABILITY

All raw datasets and homemade python full code to sort data of behavioral tests will be made available upon request.

## ACKNOWLEDGEMENTS

We thank D.Blum, L. Dupré, R. Lesourne, R. Liblau, C. Malnou, S. Marion and F. Masson for critical reading of the manuscript, F. Alt and A. Waisman for kindly agreeing to provide the floxed H2ax and Il1r1 transgenic mice, R. Balouzat, H. Bes, D. Barry, J. Bonnet, F. Chaboud, E. Debon, S. Fresse, A. J. Leblond, El Manfaloti, S. Tetot, & all services part of ANEXPLO-CREFRE for ethical care of our models, S. Allart, V. Duplan, S. Lachambre, L. Lobjois, F. L’Faqihi-Olive, V. Duplan-Eche, A.-L. Iscache, A. Chaubet, Y. Aubert & R. Romieu-Mourez from the microscopy, flow cytometry, transcriptomics and immunomonitoring core facilities of Infinity for technical help. This work was supported by institutional grants from Inserm, Marie Curie RI Europe H2020 to ES, PIA PARAFRAP Consortium (ANR-11-LABX0024 to NB), PIA ANINFIMIP equipment (ANR-11-EQPX-0003 to NB), Agence Nationale pour la Recherche (ANR-18-CE15-0015 MICCHROB to NB and ES ; ANR-22-CE14-0053 NINTENDO to ES and NB), Fondation pour la Recherche sur le Cerveau AAP2021 to NB, ANRS0366 to NB, and from Association France Alzheimer, NARSAD Brain and Behavior Foundation to ES. FHM was supported by Boehringer Ingelheim Fonds, and MB and BS by Fondation Vaincre Alzheimer. This work is part of M.B., F.S., and F.H.M. thesis projects.

## AUTHOR INFORMATION

These authors contributed equally: Marcy Belloy and Benjamin Schmitt.

These authors jointly supervised this work: Nicolas Blanchard and Elsa Suberbielle.

### Contributions

N.B, M.B., B.A.M.S, and E.S. designed preclinical experiments. D.G.D. and E.S. designed *in vitro* experiments on primary neuron cultures. M.B, , F.H.M., C.P., B.A.M.S. and E.S. performed and analysed experiments. M.B., E.B.P. and A.A. performed parasite handling and fitness analyses. M.B, R.B., R.E., F.H.M., C.P., B.A.M.S. and E.S. performed mouse husbandry and surgical procedures. A.A., M.B., E.B.P., R.B., and C.P., and E.S. processed specimens for RNA-seq and immunoprofiling and M.A., M.B., C.P., and B.A.M.S carried out bioinformatic analyses. N.B. and E.S. wrote the manuscript. All authors read, edited, and approved the final manuscript.

## ETHICAL DECLARATIONS

The authors declare no competing interests.

---

**Extended Data Figure 1:**
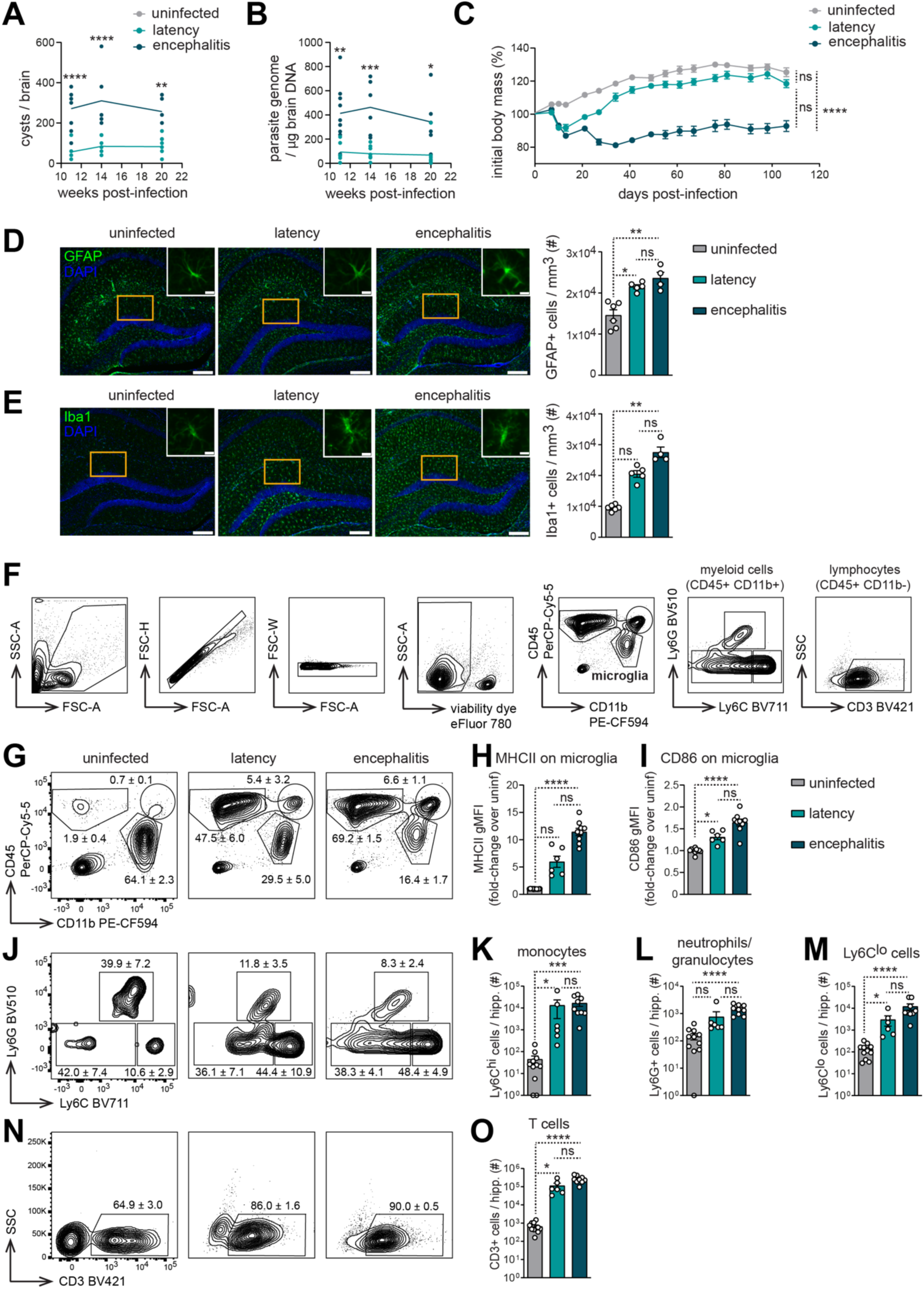
*T. gondii* infected mice display immune cell infiltration, increased astrocyte numbers and microgliosis in the hippocampus. Nine-weeks old C57BL/6J male mice were infected intraperitoneally with 200 tachyzoites of *Tg*.GRA6-OVA (latency) or *Tg*.SAG1-OVA (encephalitis) or injected with PBS as a control (uninfected). (**A**) Cyst number and (**B**) the amount of parasite genomes compared to brain mass allowed to follow the total parasite burden in the brain over time once the initial phase of brain invasion was done (mixed-effect analysis, n= 6-9 mice per group and per time point pooled from three experiments). (**C**) Percentage of initial body mass was monitored throughout infection (Kruskal-Wallis and Dunn’s multiple comparison test performed on the area under the curve, n = 7-9 mice per group from one representative experiment). (**D-E**) Cell densities of (**D**) astrocytes or (**E**) microglial cells were assessed in the dentate gyrus by immunofluorescence staining of GFAP-positive (**D**) or IBA1-positive (**E**) cells (green) with DAPI-counterstained nuclei (blue) in uninfected *vs* chronically infected mice (latency and encephalitis). Scale bar, 200 µm. Representative micrographs are shown on the left of the panel, including an orange rectangle area depicting the zone where the counting was performed. White inset displays a representative cell among the counting area. Scale bar, 10 µm. Average cell densities in dentate gyrus region of the hippocampus are shown on the right of the panel (Kruskal-Wallis, Dunn’s multiple comparison tests, n = 4-6 mice per group pooled from two independent experiments performed between 15 and 19 wpi). Each dot represents the mean value for an individual. (**F-O**) Microglia activation and immune cell infiltrates in the hippocampus were assessed by flow cytometry at 11 wpi. (**F**) The flow cytometry gating strategy is shown. (**G, J, N**) Representative contour plots with numbers indicating the mean percentage ± s.e.m. of: **G)** CD45+ CD11b-(lymphocytes), CD45+ CD11b+ (inflammatory myeloid cells) and CD45inter CD11b+ (microglia) populations out of live singlet cells, in **J**) Ly6G-Ly6C^hi^ (monocytes), Ly6C^lo^ (monocyte-derived dendritic cells and macrophages), Ly6G+ (neutrophils/granulocytes) populations out of CD45+ CD11b+ cells, and in **N**) CD3+ cells (T lymphocytes) out of CD45+ CD11b-population. (**H-I**) Markers of activation of microglia were quantified as MHC class II (**H**) and (**I**) CD86 fold-change surface expression on microglia of infected over uninfected condition. Absolute numbers of monocytes (**K**), neutrophils/granulocytes (**L**), Ly6C^lo^ cells (**M**), or T lymphocytes (**O**) were analyzed in the hippocampus. *n* = 6-11 mice per group pooled from two independent experiments. Each dot represents an individual value. ns: not significant, *p<0.05, **p<0.01, ***p<0.001, ****p<0.0001, by Kruskal-Wallis, Dunn’s multiple comparison test. Graphs display mean ± s.e.m.

**Extended Data Figure 2:**
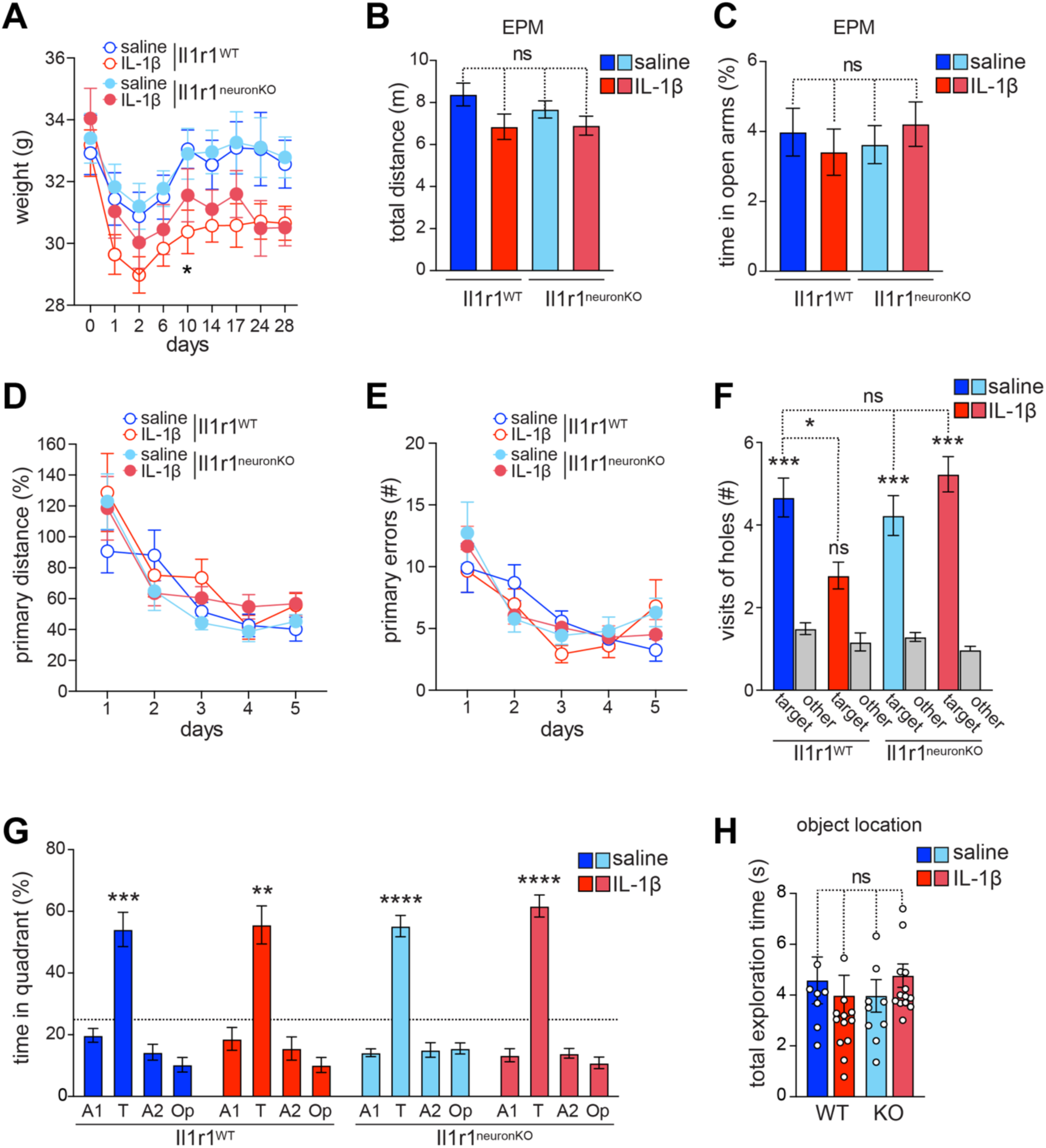
Neuronal IL-1R1 signaling in the adult hippocampus is not required for learning, consolidation nor recall of spatial memory, however is critically involved in IL-1β-induced memory impairment. *CamKII*α*-Cre-ERT2* : *Il1r1*^fl/fl^transgenic mice, in which the IL-1R1 receptor to IL-1β was knocked out in glutamatergic excitatory neurons of the forebrain (*Cre^+^*, *Il1r1*^neuronKO^) upon tamoxifen treatment and their *Cre*^-^ *Il1r1*^WT^ littermates were implanted with subcutaneous osmotic mini-pumps containing saline or IL-1β diluted in saline solution (5 µg/kg/day) at 3,5-4-5 months. (**A**) Body mass was monitored throughout the experiments post implantation (Impact of the time (p<0.0001) and the interaction of the time with the group by mixed effects model analysis (REML) (*p<0.05 by Dunn’s test), *n* = 16-20 mice per group pooled from three independent experiments. (**B-H**) Starting 3 weeks post-implantation, mice were tested in a series of behavioral tests: the elevated plus maze (EPM, B-C), the Barnes maze (**D-G**), and the object location task (**H**). (**B-C**) in the EPM, all mice behaved similarly in terms of distance run during the first 5 minutes (**B**) or anxiety levels as shown by the percentage of exploration time spent within the open arms (**C**) (*n* = 16-26 mice per group of treatment pooled from 3 independent cohorts and results analyzed by Bonferroni post-hoc tests). (**D-G**) In the Barnes maze, learning curves show mean daily distance run as a percentage of the mean distance run during the first trial by the mice of the saline-treated *Il1r1*^WT^ group for each cohort (**D**), or mean numbers of errors of holes visits (**E**) prior to find the target hole connected to a hidden exit box. Effect of the time (p<0.0001) was analyzed by repeated measures mixed-effects models. During a 90-s probe trial run 24 h after the last training trial in the maze, remote memory was assessed. (**F**) the numbers of visits of the original exit hole (target) *vs*. other holes in other quadrants (other) during the probe trial were counted. Results were analyzed by paired *t*-tests between target and other or by Dunnett’s post-hoc tests compared to the saline group as indicated by brackets. (**G**) Percentage of time spent by mice in the target quadrant *vs*. the three other quadrants during the probe trial. Results were tested by one-sample t-test compared to chance (25%). n = 10–13 mice per group of treatment pooled from 2 independent cohorts. (**H**) Bars represent total time of exploration of both object in the object location test shown in Figure 3. n = 8–14 mice per group of treatment pooled from 2 independent cohorts. Each dot represents the performance of one mouse. ns, non-significant, *p<0.05, **p<0.01, ***p<0.001, ****p<0.0001. Graphs show means ± s.e.m.

**Extended Data Figure 3:**
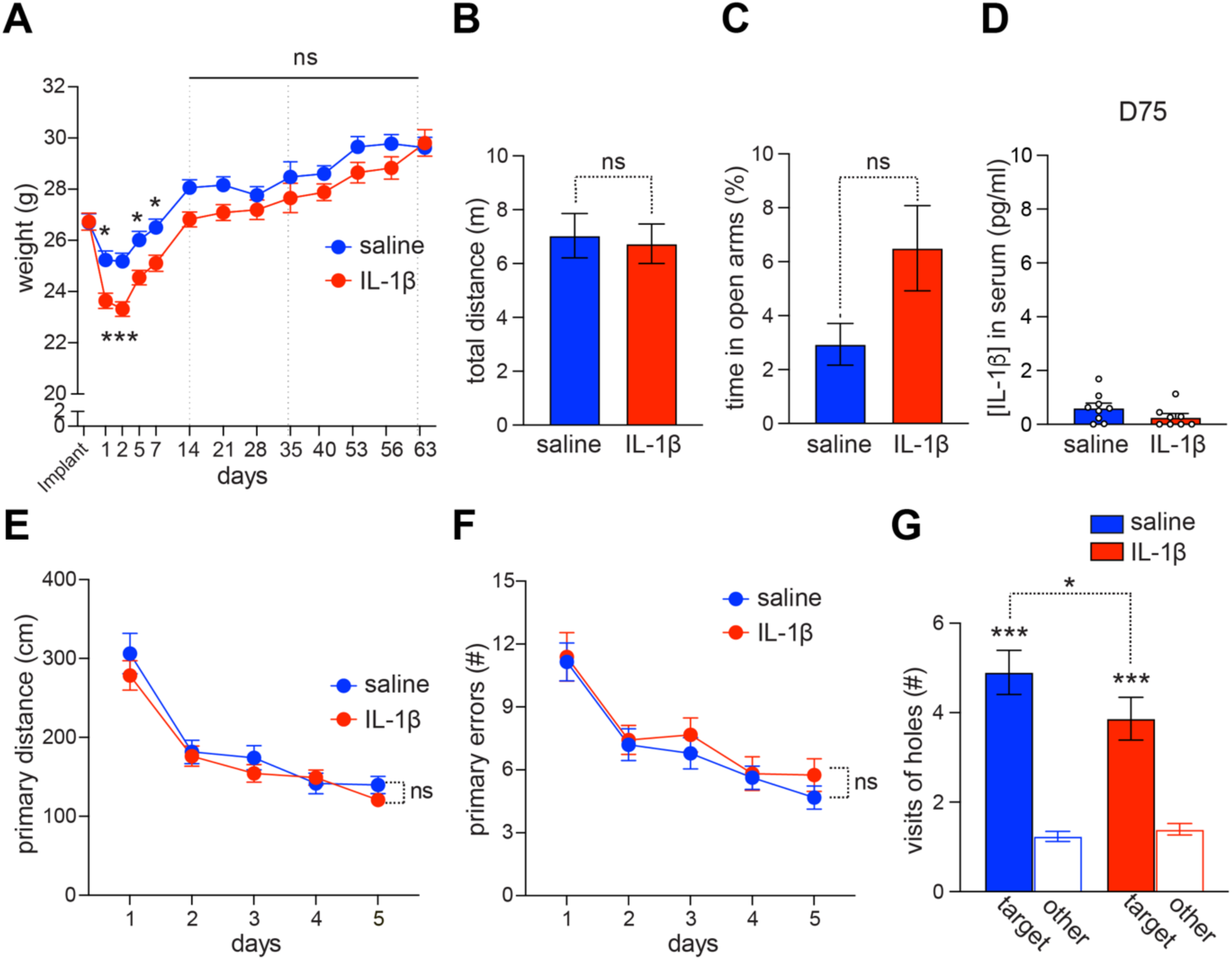
Chronic IL-1β does not impact anxiety or activity but causes long-lasting memory impairment despite clearance of the cytokine from the body. C57BL/6J male mice were subcutaneously implanted with osmotic mini-pumps containing saline or IL-1β diluted in saline solution (5 µg/kg/day) for 35 days. (**A**) Body mass was monitored throughout the experiments post implantation (Impact of the time (p<0.0001), treatment (p=0.03) and the interaction of both (p<0.0001) analyzed by mixed effects model analysis (REML), *p<0.05, ***p<0.001 by Bonferroni post-hoc tests, *n* = 29 mice per group pooled from three independent experiments. 3 weeks post-implantation, mice were tested in the EPM (**B-C**), and 1 month later, in the (Barnes maze (**E-G**). (**B-C**) in the EPM, all mice behaved similarly in terms of distance run during the first 5 minutes (**B**) or anxiety levels as shown by the percentages of exploration time spent within the open arms (**C**) (*n* = 16 mice per group of treatment pooled from 2 independent cohorts and results analyzed by Welch’s t-tests). (**D**) IL-1β concentrations in the serum were assessed by ELISA 3 days after the Barnes maze probe test in saline controls or IL-1β-treated group of mice. n= 8-9 mice per group from one experiment (unpaired Student’s *t* test). Each dot represents the value for one individual. (**E-G**) In the Barnes maze, learning curves show mean daily distance run (**E**), or mean numbers of errors of holes visits (**F**) prior to find the target hole connected to a hidden exit box. Effect of the time (p<0.0001) was analyzed by repeated measures mixed-effects models. ns, non-significant by Dunn’s tests (**G**) Number of visits of the original exit hole (target) *vs*. other holes in other quadrants (other) during the probe trial. Results were analyzed by paired *t*-tests between target and other or by Student-t tests compared to the saline group as indicated by brackets. *n* = 14–20 mice per group of treatment pooled from 2 independent cohorts. ns, non-significant, *p<0.05, **p<0.01, ***p<0.001. Values are indicated as means ± s.e.m.

**Extended Data Figure 4:**
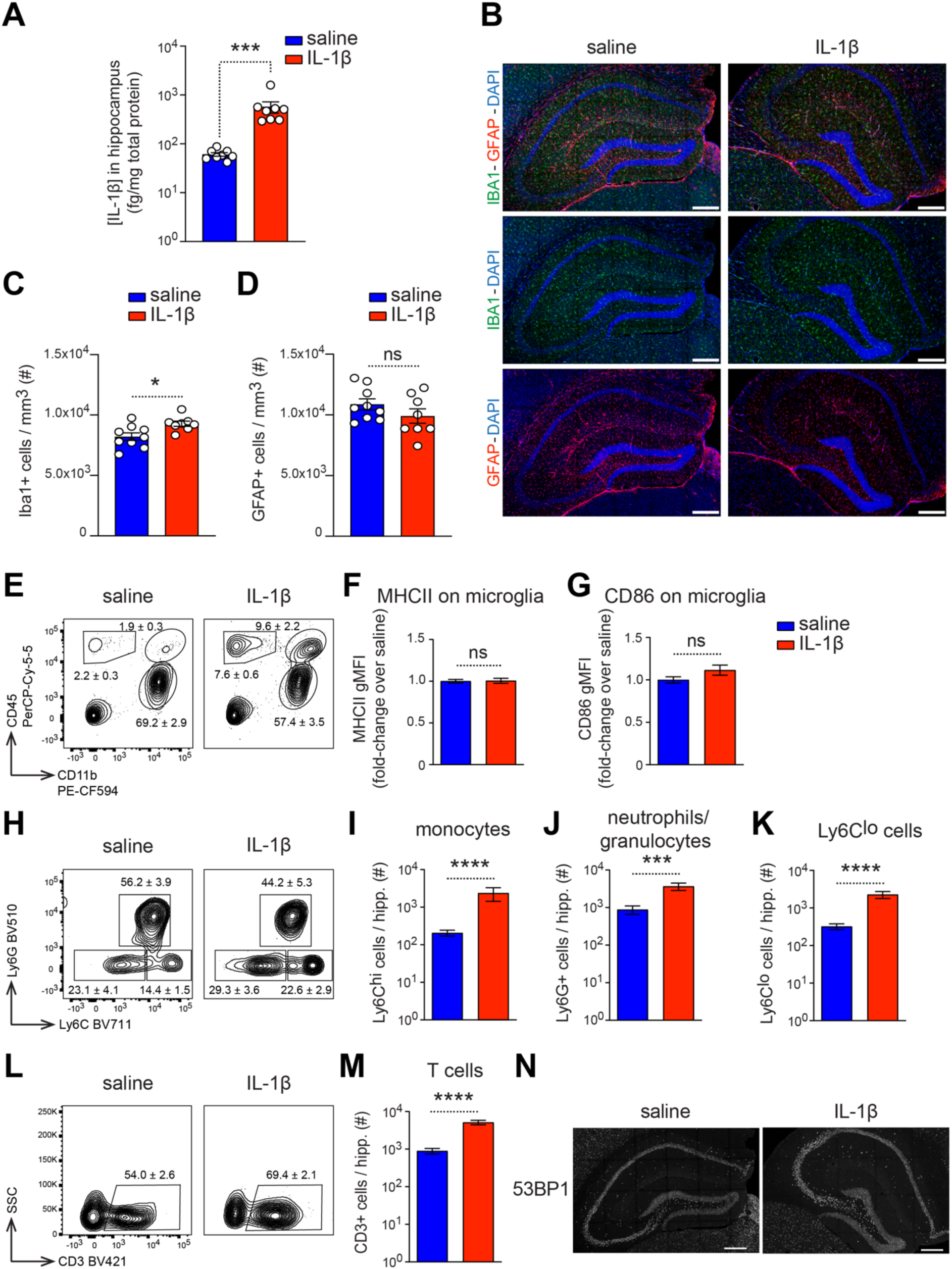
Chronic exposure to IL-1β induces peripheral immune cell recruitment in the hippocampus without overt microglia activation. C57BL/6J male mice were subcutaneously implanted with osmotic mini-pumps containing saline or IL-1β diluted in saline solution (5 µg/kg/day) for 27 days. **A**) IL-1β concentration was measured in the hippocampus by Meso Scale Discovery Electrochemiluminescence (Mann-Whitney test, n = 8 mice per group pooled from 2 independent cohorts). (**B-D**) Cell densities of (**B, C**) microglial cells or (**B, D**) astrocytes were assessed in the dentate gyrus by immunofluorescence stainings (**A**) of IBA1-positive cells (green), GFAP-positive cells (red) with DAPI-counterstained nuclei (blue) in coronal sections from the mice. (**A**) Representative micrographs are shown. The counting area is similarly positioned as in Extended Data Figure 1D. Average cell densities were analyzed by Student t tests, (n = 9 mice per group from one experiment of C57BL6/J mice). Each dot represents an individual value. (**E-M**) Microglia activation and immune cell infiltrates in the hippocampus were assessed by flow cytometry. (**E, H, L**) Representative contour plots with numbers indicating the mean percentage ± sem of (**E)** CD45+ CD11b-(lymphocytes), CD45+ CD11b+ (inflammatory myeloid cells) and CD45inter CD11b+ (microglia) populations out of live singlet cells, in (**H**) Ly6G-Ly6C^hi^ (monocytes), Ly6C^lo^ (monocyte-derived dendritic cells and macrophages), Ly6G+ (neutrophils/granulocytes) populations out of CD45+ CD11b+ cells, in (**L**) CD3+ cells (T lymphocytes) out of CD45+ CD11b-population. (**F**) MHC class II and (**G**) CD86 fold-change surface expression on microglia in IL-1β over saline condition (unpaired Student’s t tests). Absolute numbers of (**I**) monocytes, (**J**) neutrophils/granulocytes, (**K**) Ly6C^lo^ cells and, (**M**) T lymphocytes (Mann-Whitney test). (**N**) Confocal micrographs of the hippocampus of two mice stained for DSB (53BP1, gray), indicating no changes in morphology and representative specific diffuse staining of 53BP1 in neurons. Scale bar, 200 μm. n = 10 to 14 mice per group pooled from two independent experiments. ns, non-significant, ***p<0.001, ****p≤0.0001. Graphs show means ± s.e.m.

**Extended Data Figure 5:**
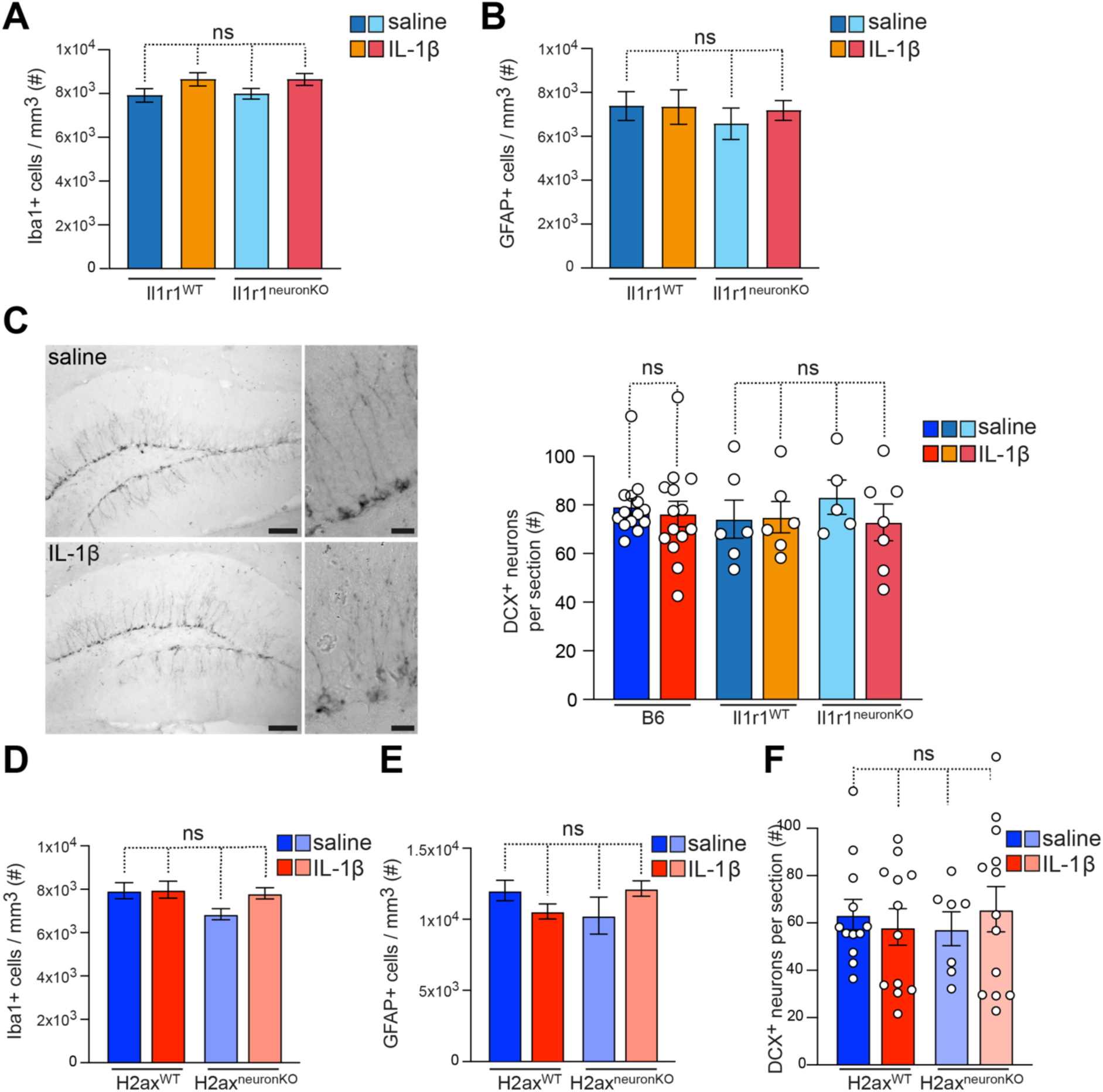
Astrocytes, microglia nor new-born neurons cell numbers remain unaffected in *Il1r1*^neuronKO^ nor *H2ax*^neuronKO^ mice despite chronic systemic exposure to low levels of IL-1β. Cell densities of (**A, D**) microglial cells or (**B, E**) astrocytes were assessed in the dentate gyrus by immunofluorescence stainings (**A**) of IBA1-positive cells (green), GFAP-positive cells (red) with DAPI-counterstained nuclei (blue) in coronal sections from transgenic mice, in which the *Il1r1* receptor to IL-1β was knocked out in neurons (*Il1r1*^neuronKO^ or not (for Cre^-^ (*Il1r1*^WT^) littermates) upon tamoxifen treatment (**A-B**), or in transgenic mice, in which *H2ax* was knocked out in neurons (*H2ax*^neuronKO^ or not (for Cre^-^ (*H2ax*^WT^) littermates) upon tamoxifen treatment (**D-E**) and after 28 days of infusion of saline versus IL-1β. The counting area is similarly positioned as in Extended Data Figure 1D. Average cell densities were analyzed by Student t tests, (n=11-17 mice per group of *Il1r1* transgenic mice pooled from two independent experiments, and n=11-23 mice per group of *H2ax* transgenic mice pooled from three independent experiments). (**C & F**) The number of newly born granule cells in the DG was determined by immunostaining of coronal brain sections for doublecortin (DCX) and counting immunoreactive neurons in the DG regions of three sections per mouse. (**C**) Representative micrographs (Scale bar, 100 µm) with details showed in right insets (Scale bar, 20 µm). On the right, the average number of immunoreactive neurons per section is shown (*n* = 14 mice per group pooled from two independent experiments of C57BL6/J mice and n=5-7 mice per group of *Il1r1* transgenic mice from one experiment). (**F**) *n* = 7-13 mice per group *H2ax* transgenic mice pooled from two independent experiments. ns, non significant, by Bonferroni post-hoc tests. Each dot represents the mean value for an individual. Values are shown as means ± s.e.m.

## Notes

### Competing Interest Statement

The authors have declared no competing interest.

